# Harnessing systemic glycolysis-TCA cycle axis to boost the host defense against newborn infection

**DOI:** 10.1101/2025.04.23.650144

**Authors:** Ziyuan Wu, Nguyen Tran Nam Tien, Björn Klabunde, Karoline Aasmul-Olsen, Simone Margaard Offersen, Nguyen Thi Hai Yen, Tik Muk, Anna Hammerich Thysen, Susanne Brix, Nicklas Brustad, Tingting Wang, Jakob Stokholm, Klaus Bønnelykke, Anders Brunse, Nguyen Phuoc Long, Bo Chawes, Ole Bæk, Duc Ninh Nguyen

**Affiliations:** Comparative Pediatrics, Department of Veterinary and Animal Sciences, University of Copenhagen, Frederiksberg, Denmark; Department of Pharmacology and PharmacoGenomics Research Center, Inje University College of Medicine, Busan 47392, Republic of Korea; Department of Biotechnology and Biomedicine, Technical University of Denmark, Lyngby, Denmark; COPSAC, Copenhagen Prospective Studies on Asthma in Childhood, Herlev and Gentofte Hospital, University of Copenhagen, Copenhagen, Denmark; Department of Clinical Medicine, Faculty of Health and Medical Sciences, University of Copenhagen, Copenhagen, Denmark; BRIDGE Translational Excellence Programme, Faculty of Health Sciences, University of Copenhagen, Copenhagen, Denmark

## Abstract

Energy metabolism and immune response are tightly connected, but it is poorly understood how this interplay is regulated in early life to dictate host defense strategy, infection risks and severity. This interplay is particularly relevant for preterm, low birthweight or otherwise immunocompromised infants, who have poor metabolic control and increased risks of sepsis. Here, we utilized data from the COPSAC_2010_ cohort with 700 mother-child pairs and showed that plasma levels of TCA cycle metabolites in early life were associated with reduced childhood risk of bacterial infection and an attenuated systemic inflammatory response. Next, we explored how two distinct nutritional strategies, which were aimed at boosting TCA cycle activity instead of glycolysis, impacted neonatal host defense against a serious bloodstream infection in preterm piglets. Substituting galactose for glucose in parenteral nutrition enhanced disease tolerance in early phase of infection and overall glucose homeostasis, improving survival. Further, combining glucose restriction with supplementation of glucogenic amino acids conferred glycemic control and completely prevented sepsis and abnormal changes of organ injury markers. Mechanistically, this intervention enhanced both disease resistance and tolerance, accompanied by metabolic rewiring from glycolysis towards gluconeogenesis, TCA cycle activity and oxidative phosphorylation. Thus, optimized nutritional strategies controlling the interplay of energy metabolism and host defense may be lifesaving for infected infants.

**In brief:** Newborns rely on two distinct defense strategies to combat infections in early life. Disease resistance, fueled by aerobic glycolysis, seeks to actively eliminate microorganisms while disease tolerance, fueled by mitochondrial oxidative phosphorylation, seeks to reduce collateral tissue damage during infections. We found that in healthy human newborns, increased plasma levels of metabolites from the tricarboxylic acid (TCA) cycle were associated with lower burden of childhood infections and reduced pro-inflammatory status. In a newborn animal model of bloodstream infection, nutritional strategies boosting systemic TCA cycle activity, while reducing aerobic glycolysis, enhanced both host disease tolerance and resistance, thereby improving survival. These findings could pave a path for improved infection management in human newborns.

**Highlights:**

- In healthy children, higher plasma levels of TCA cycle metabolites are associated with lower infection risks and systemic inflammation.
- In a neonatal infection model, the supply of galactose, instead of glucose, improves host glucose homeostasis and TCA cycle activity, improving disease tolerance and survival.
- A combination of glucose restriction and glucogenic amino acid supply also improves TCA cycle activity, enhancing both disease resistance and tolerance and completely preventing lethal sepsis.

## Introduction

Newborn infants are highly susceptible to infections that can progress to neonatal sepsis, a state of excessive inflammation causing organ damage with high risks of mortality^1^. This infection risk in newborns has conventionally been attributed to their immature immune system with limited capacity to respond to exogenous stimuli^2^. However, this hypo-responsiveness may also be influenced by the newborn energy metabolism^3^. In adults with sufficient energy stores, immune cells undergo a metabolic shift from oxidative phosphorylation (OxPhos) to glycolysis upon infection to mediate fast ATP production, fueling pro-inflammatory responses to eliminate invading pathogens at a cost to host fitness, a defense strategy termed disease resistance^4^. Newborns in contrast, especially those born prematurely or at low birthweight, have limited energy reserves, but high energy demands for growth. As a result, they are programmed to prioritize their energy for vital organ functions rather than for inflammatory responses to infection, leading to the tendency to tolerate invading pathogens, a defense strategy termed disease tolerance^3,4^. However, when pathogen growth exceeds a certain threshold, the glycolysis-mediated resistance mechanisms occur in an excessive manner, causing collateral tissue damage and organ dysfunctions^5^.

We have shown that boosting of disease tolerance phenotype in infected preterm pigs is accompanied to increased survival and enhanced hepatic OxPhos^5^. In addition, a reduction of parenteral glucose intake during serious bloodstream infections improves survival, via dampened glycolysis and increased OxPhos activity in the liver, resulting in more balanced immune responses^6,7^. A key component of mitochondrial OxPhos is the tricarboxylic acid (TCA) cycle, a cyclic metabolic pathway that can be fueled by various intermediary metabolites such as amino acids, acetyl coenzyme A and pyruvate. Through a series of enzymatic reactions, the TCA cycle generates electron carriers to facilitate the electron transport chain, for adenosine triphosphate (ATP) production. However, TCA cycle metabolites are not confined to mitochondria but are actively transported to the cytosol, nucleus and extracellular space where they can serve as signaling molecules^8^. In macrophages, accumulation of both α-ketoglutarate and succinate enhances the M2 anti-inflammatory phenotype pathway ^9,10^, while these molecules together with itaconate elevate the cell-responding threshold to lipopolysaccharide (LPS), an effect mirroring disease tolerance ^9–11^. In contrast, depletion of α-ketoglutarate increases the severity of LPS-induced sepsis in adult mice ^12^. These effects suggest that the overall activity of the TCA cycle may affect disease tolerance against serious infections. Importantly, activity of the TCA cycle also generates the necessary metabolites for gluconeogenesis which, in a reverse set of reactions to glycolysis, leads to the *de novo* formation of glucose^8^. In line with this, we have previously shown that reduction in glucose intake increases hepatic gluconeogenesis and infection survival in newborn preterm pigs^7^. As such, enhancement of the activity of the TCA cycle may help maintain normoglycemia upon limited glucose intake.

In the current study, we first examined the association between plasma levels of TCA cycle metabolites and infection risks and immune function during early life, using available data from a cohort of 700 mother-child pairs^13^. Thereafter, combining studies using a macrophage cell line and a preterm pig model of newborn bacterial bloodstream infection, we explored the capacity of nutritional strategies to optimize the host infection defense by boosting activities of the TCA cycle instead of glycolysis. We found that increased levels of key TCA cycle metabolites at birth were associated with reduced infection risks during the first 3 years of life. Importantly, during childhood, these metabolites were also associated with anti-inflammatory responses to exogenous antigens of blood immune cells. Next, we found that nourishing preterm piglets with galactose instead of glucose improved blood glucose homeostasis and disease tolerance against infection, whereas replacing a major fraction of glucose intake with glucogenic amino acids simultaneously enhanced disease resistance and tolerance, in turn maximizing infection survival. The clinical benefits of both these strategies were associated with hepatic metabolic rewiring to increased TCA cycle activity and reduced levels of organ injury markers. By combining data from a large human birth cohort with *in vitro* and *in vivo* experiments, our study provides both mechanistic insights, proof-of-concept and translational value for the future development of optimized nutritional approaches in infected infants.

## Results

### Mother-child cohort characterization

To investigate the relationship between TCA cycle metabolites, infection risk, and immune function in human newborns, we analyzed data from the Copenhagen Prospective Studies on Asthma in Childhood 2010 (COPSAC_2010_) birth cohort. The COPSAC_2010_ is a population-based cohort consisting of 700 mother-child pairs, who were closely monitored from birth through the first three years of life and onwards with daily dairy registration of infections at age 0-3 years^14^, combined with longitudinal biosampling from multiple compartments including blood for immune function and omics assessments^13,15^. Detailed characteristics of the cohort has been published previously^16,17^. The comprehensive documentation of common childhood infections was combined with plasma LC-MS metabolome and immune profiling to provide a detailed dataset for analysis (**Figure 1A**). Plasma metabolome data was available from the mother at one week postpartum (n=678), and from the children at 6 (n=562) and 18 months (n=538). At all the timepoints, semi-quantitative levels of citrate, α-ketoglutarate, succinate, fumarate and malate could be assessed within quality control limits.

**Figure 1.**
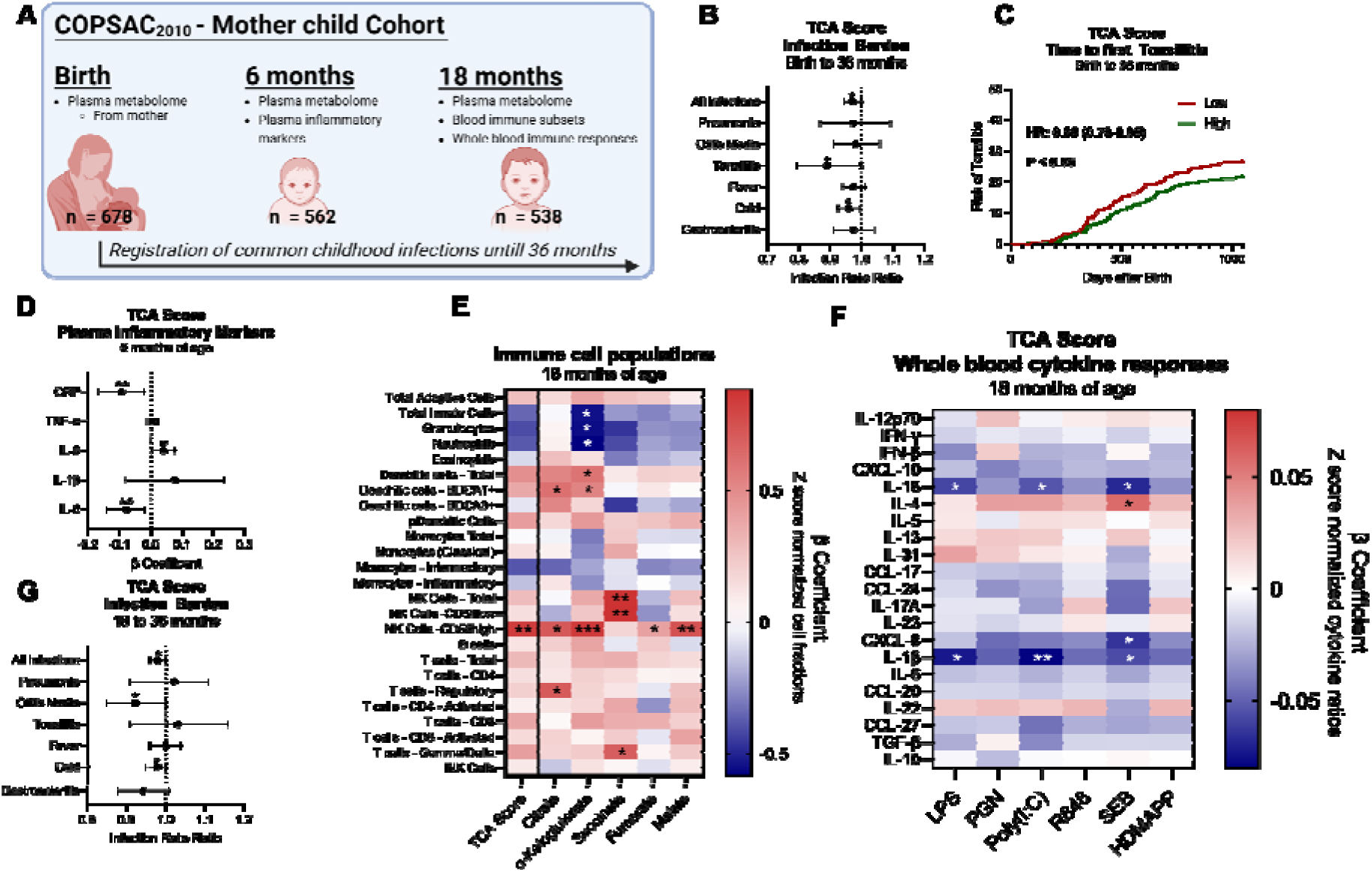
Plasma levels of TCA cycle metabolites are associated with infection burden and immune responses in healthy young children. **(A)** Schematic overview of the cohort. Plasma metabolomics samples were obtained from mothers at birth and children at 6 and 18 months. Levels of plasma TCA cycle metabolites were associated with the burden of common infections until 36 months of age, plasma inflammatory markers (6 months), and immune cell subsets and cytokine responses (18 months). **(B)** Effect of maternal TCA scores on infection counts from birth to 36 months of age, shown as infection risk ratios with corresponding 95% confidenc intervals. **(C)** Effect of maternal TCA scores on time to first tonsillitis infection, shown as Kaplan Meier curves of children with higher (green) or lower (red) TCA scores than the median. The results of Cox proportional hazard models are shown as text. **(D)** Association between TCA scores at 6 months of age and plasma inflammatory markers CRP: C-reactive protein, TNF: Tumor necrosis factor, IL: Interleukin. Shown as β-coefficients with corresponding 95% confidence intervals. **(E)** Heatmap showing the association between TCA score and individual TCA metabolites at 18 months of age and zscore normalized fractions of immune cell determined by flow cytometry. Shown as β-coefficients where red color indicates positive, and blue has a negative association. **(F)** Heatmap showing the association between TCA score at 18 months of age and zscore normalized cytokine ratios following whole blood stimulation with six pattern recognition receptor agonists. Shown as β-coefficients where red color indicates positive, and blue has a negative association. LPS: Lipopolysaccharide, PGN: Peptidoglycan, R848: Imidazoquinoline, SEB Staphylococcal enterotoxin B, HDMAPP: 1-Hydroxy-2-methyl-2-buten-4-yl 4-diphosphate. **(G)** Association between PC_1_ scores at 18 months of age and infection counts at 36 months of age is shown as infection risk ratios with corresponding 95% confidenc intervals. *P-value < 0.05, **P-value < 0.01, and ***P-value < 0.001. Panel A was created using Biorender.com.

Of the 678 children included in the infection burden analysis, all but six experienced at least one infection episode within their first three years, with a median of 16 ± 8 episodes. Infections were categorized as suspected bacterial (e.g., pneumonia, acute otitis media, tonsilitis) or suspected viral (e.g., cold symptoms, gastroenteritis), while episodes of fever without localized symptoms were also recorded. At 6 months of age, systemic inflammation markers in plasma were measured in all 562 infants, while at 18 months, immune function was assessed in 495 of 538 children using flow cytometry and whole blood stimulations with six different pattern recognition receptor (PRR) agonists or T cell ligands (**Figure 1A**).

### TCA cycle metabolites, infection burden and inflammatory responses in human newborns

The five key annotated TCA cycle metabolites (α-ketoglutarate, malate, succinate, citrate and fumarate) were identified from plasma metabolome data with identity verified using authentic chemical standards. These metabolites were pre-selected for an unsupervised principal component analysis (PCA) to obtain components representing overall TCA cycle activity. In both the maternal metabolome at birth and the child metabolome at 6 and 18 months of age, individual TCA cycle metabolites contributed nearly equally and positively to Principal Component 1 (PC, from now assigned as TCA score), which explained 47-49% of the variation at each timepoint (**Figure S1A-F**).

When correcting for relevant confounders, higher maternal TCA cycle activity at birth, as reflected by higher TCA score, was associated with a reduced overall infection count, driven by fewer episodes of tonsilitis and cold symptoms in children during the first 3 years of life, as estimated by quasi-poisson regression (**Figure 1B**). Higher TCA scores were also associated with increased time to first tonsilitis episode, as estimated by a cox proportional hazard model (**Figure 1C**), but not for other infection types. These effects appeared to be driven primarily by higher levels of citrate, fumarate and malate (**Table S1**). We have previously shown a high correlation between the maternal and newborn plasma metabolome, making the maternal metabolome at birth suitable for our analysis^18^.

At 6 months of age, higher child TCA scores were associated with lower plasma levels of Interleukin 6 (IL-6) and C-reactive protein, but higher levels of IL-8 (**Figure 1D**). At 18 months of age, higher child TCA scores were associated with higher frequencies of CD56^Bright^ Natural Killer (NK) cells (**Figure 1E**). Individually, α-ketoglutarate by itself correlated with lower frequencies of neutrophils and higher frequencies of BDCA-1^+^ dendritic cells (DCs, **Figure 1E**). Furthermore, citrate and succinate correlated with higher frequencies of regulatory T cells (T_Reg_) and γδ-T cells respectively (**Figure 1E**). These findings align with known metabolic dependencies of immune cell subsets as neutrophils predominantly relying on glycolysis, while CD56^Bright^ NK cells, DCs, T_Reg_, and γδ T cells depend on OxPhos^19,20^.

The changes in immune cell subsets at 18 months of age were reflected in cytokine responses to various PRR agonists and T cell ligands. Higher TCA scores were associated with reduced responses of the pro-inflammatory cytokines IL-1β and IL-18 after stimulation of whole blood with LPS, Poly(I:C) and SEB and increased IL-4 responses to SEB (**Figure 1F**). Among the five TCA cycle metabolites, levels of citrate were consistently associated with increased levels of the anti-inflammatory cytokine interleukin-10 (IL-10) across 5 of the 6 different PRR agonists stimulations (**Figure S1G**). Likewise, α-ketoglutarate, succinate, fumarate and malate were associated with reduced responses of several pro-inflammatory cytokines (CXCL-10, IL-18, CXCL-8 and IL-1β) to several ligands (**Figure S1, H-K**). Importantly, TCA scores at 18 months were still associated with a lower risk of total infection episodes, now driven by fewer episodes of acute otitis media and cold symptoms, in the period between 18-36 months of age (**Figure 1G**).

Together, these results suggest that plasma levels of TCA cycle metabolites at birth and during infancy correlate with differences in infection burdens, systemic inflammation, immune cell subsets, and immune function in healthy human newborns. Even though plasma metabolites reflect the whole host body metabolism, not specifically immune cell metabolism, these findings underscore the importance of energy metabolic pathways in shaping newborn immune responses.

Likewise, there may be differences in each of the TCA metabolites, as e.g. citrate consistently associated with increases in anti-inflammatory cell types and cytokine responses, whereas α-ketoglutarate associated with reductions in innate cell populations and pro-inflammatory cytokine responses. These associations may be relevant during serious infections in young children. Using a neonatal sepsis model in infected preterm pigs, we have shown that infected survivors consistently had higher levels of TCA cycle plasma metabolites, relative to infected non-survivors, underscoring the potential role of TCA cycle activity in regulating host defense^5^. Therefore, building on these insights, we aimed to explore the causal impacts of modulated TCA cycle activity by novel nutritional interventions on host defense during neonatal infections.

### Supply of galactose instead of glucose improves disease tolerance and infection survival

With the sepsis model in infected preterm piglets, we have demonstrated that a standard glucose supply, similar to that given to preterm infants, results in hyperglycemia, excessive glycolysis-induced inflammation and sepsis, while glucose restriction prevents sepsis but induces hypoglycemia^6,7^. As such, here we first aimed to explore how an alternative carbohydrate supply, with identical energy density, could reduce the rate of glycolysis and improve TCA cycle activity during newborn infections. Galactose is a monosaccharide bound to glucose in the natural structure of lactose, the main carbohydrate of milk and cleaved by lactase during gastrointestinal digestion prior to absorption. Despite its obvious safety, direct use of galactose as nutrition for infants has not been reported. Galactose metabolism starts with the Leloir pathway prior to entering glycolysis^21^, and we hypothesized that the slow conversion of galactose (compared to glucose) into glucose-6-phosphate can reduce systemic glycolytic activity during neonatal infection, increasing reliance on mitochondrial OxPhos, in turn promoting host disease tolerance with reduced inflammation.

First, we tested the hypothesis *in vitro* by culturing human macrophage-like THP1 cells in medium containing different levels of galactose and glucose, exposed to live *Staphylococcus epidermidis*, the most common pathogen detected in infected preterm infants^22^. After 6 hours of stimulation, low doses of galactose addition (10mM) reduced production of TNF-α and tended to increase that of IL-10 (**Figure 2A and B**). There was however no difference in the capability of cells to produce ATP between galactose and glucose, or between doses (**Figure 2C**) and minimal effects on gene expression (**Figure S2A**). This suggests that supplying galactose, instead of glucose, at relevant concentrations into the human bloodstream may reduce pro-inflammatory drive of immune cells without impacting their energy metabolism.

**Figure 2.**
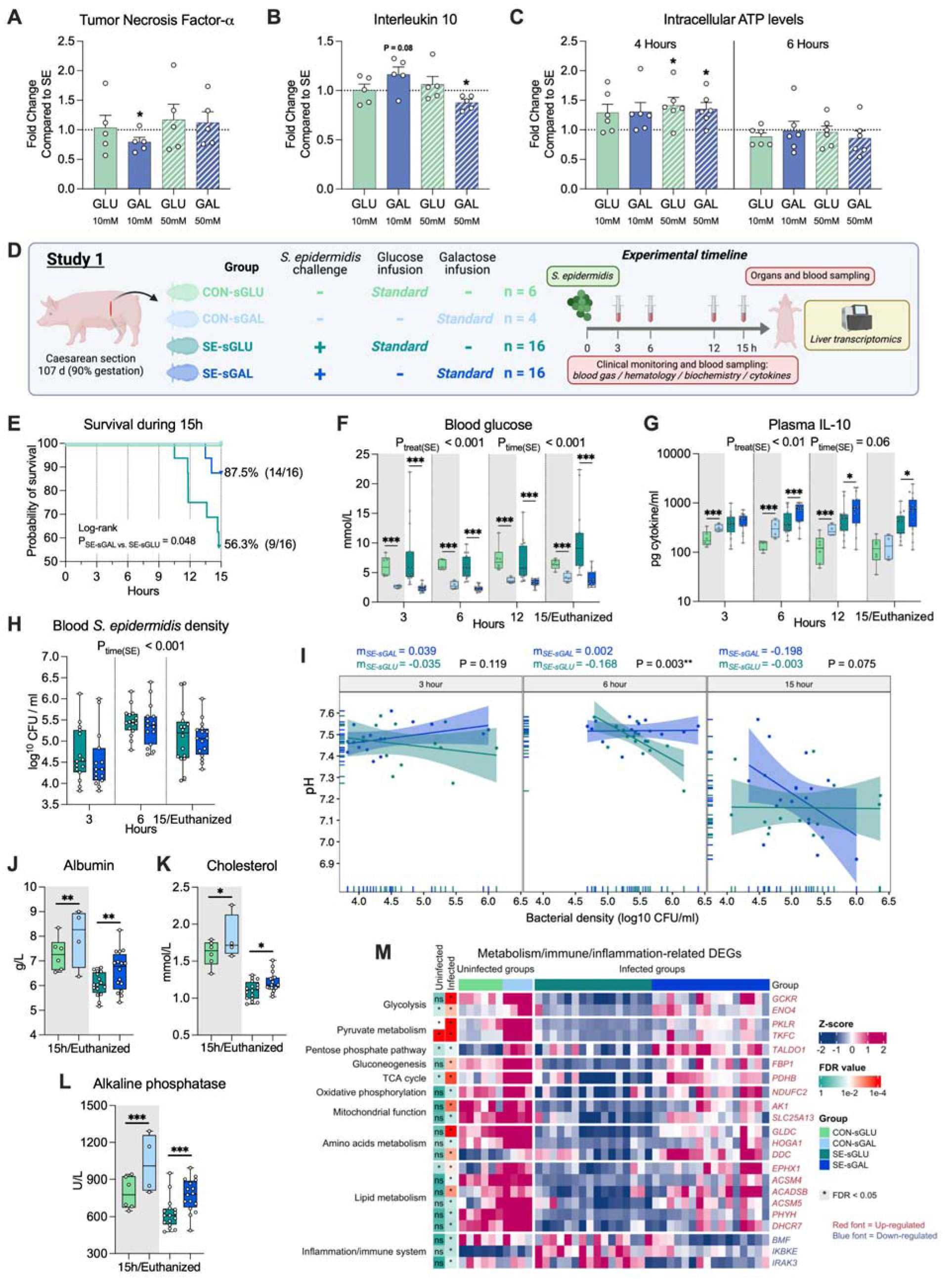
Supply of galactose instead of glucose maintains normoglycemia, improves host disease tolerance, and reduces the risk of sepsis. **(A–C)** *In vitro* human macrophage-like THP1 cells exposed to live *S. epidermidis*. Levels of tumor necrosis factor-α, interleukin 10, and intracellular ATP are shown as relative fold changes in relation to *S. epidermidis* positive control. **(D)** Schematic overview of the experimental design. Forty-two preterm piglets delivered by cesarean section were randomly assigned to groups exclusively nourished with parenteral nutrition containing either a standard glucose solution (sGLU; 10% glucose, 14.4 g/kg/d) or an equivalent concentration of galactose (sGAL, 10% galactose, 14.4 g/kg/d). Piglets then received either live *S. epidermidis* (SE, 10^9^ CFU/kg, n=32) or saline control (CON, n = 10). This setup allowed for the division of piglets into four groups: CON-sGLU (n = 6), CON-sGAL (n = 4), SE-sGLU (n = 16), and SE-sGAL (n = 16). **(E)** Survival curves of the animals throughout the experiment, displayed as the time until 15 h post-inoculation or humane euthanasia based on predefined humane endpoints. **(F–G)** Blood pH and plasma IL-10) at 3, 6, 12, and 15 h post-inoculation or at euthanasia. **(H)** Density of *S. epidermidis* in jugular venous blood samples from infected animals at 3 and 6 h or heart puncture at euthanasia. **(I)** Reaction norm analysis, performed using blood pH (as a readout for health) at 3, 6, and 15 hours, and plotted against the pathogen burdens at the same time point by linear regression. Extra sum-of-squares F Test was used to compare slopes. **(J–L)** Serum biochemistry parameters (albumin, cholesterol, and alkaline phosphatase) at 15 h post-inoculation or at euthanasia. **(M)** Heatmap illustrating top DEGs between SE-sGAL and SE-sGLU groups. **Statistics**: **(E)** A log-rank test was used to compare survival between the SE-sGAL and SE-sGLU groups. **(F–H & J–L)** Data at each time point were analyzed using a linear mixed-effects model, incorporating group, gender, and birth weight as fixed factors and litter as a random factor. *P-value < 0.05, **P-value < 0.01, and ***P-value < 0.001, compared between SE-sGAL and SE-sGLU groups at the same time point. **(F–H)** Another linear mixed-effects model was employed to probe further disparities spanning the entire experimental duration, incorporating group, time, their interaction, gender, and birth weight as fixed factors, with litter and pig ID as random factors. P_treat(SE)_, P_time(SE)_, and P_int(SE)_ denote probability values for group effect (SE-sGAL and SE-sGLU) over time, time effects, and the interaction effects between time and group in the linear mixed effects interaction model, respectively. Uninfected animals (CON) served as a reference group and were not compared directly with infected animals (SE). Statistical significance was defined as P-value < 0.05. All data were presented as box and whisker plots showing the range from minimum to maximum values. Panel D was created using Biorender.com.

Next, we tested the substitution of galactose for glucose in parenteral nutrition, using our well-established neonatal sepsis model^5–7,23^. Preterm piglets were delivered at 90% gestation, infected with *S. epidermidis* (SE), reared with standard dose (14.4g/kg/day) of glucose (sGLU, similar to what preterm infants receive) or galactose (sGAL), and intensively monitored until 15 h post-inoculation (**Figure 2D**). Compared to SE-sGLU animals, SE-sGAL animals showed lower mortality due to sepsis (**Figure 2E**), which was predefined as arterial blood pH of ≤ 7.1, together with deep lethargy, apnea or hypoperfusion. This was accompanied by a tendency of higher blood pH and lower carbon dioxide pressure (pCO_2_) in SE-sGAL animals over time (**Figure S2B-C**), suggesting that galactose supply prevented respiratory acidosis. During the first 6h, galactose-supplemented animals also showed a slight elevation of lactate (**Figure S2D**), potentially indicating a stronger glycolysis-related immune response to resist the infection. Importantly, glucose-supplemented animals showed high blood glucose levels, while both infected and uninfected galactose supplied animals were normoglycemic, with gradually increasing blood glucose levels from 2.4 to around 4 mM over time, indicating the ability to continuously convert exogenous galactose into glucose (**Figure 2F)**.

In SE-sGAL animals, plasma IL-10 was elevated (**Figure 2G**), while bacterial clearance (**Figure 2H**) and other inflammatory cytokines TNF-α and IL-6 (**Figure S2E and F**) were unchanged. However, SE-sGAL animals had less depletion of blood neutrophils, monocytes, and hemoglobin (**Figure S2G–I**). Thus, it appears that the clinical benefit of galactose supply was derived from improved disease tolerance instead of resistance, prioritizing the maintenance of normal organ function over a strong immune response to clear the pathogens. To confirm the disease tolerance phenotype, we employed a reaction norm analysis by plotting changes in host health against pathogen burdens^5^. Here, improved disease tolerance would be indicated by a shallower slope, showing the capacity to maintain health despite increasing pathogen burdens. Blood pH was used as a readout for general health, and reaction norms were generated at 3, 6, and 15 h post-inoculation (**Figure 2I**). Both infected groups displayed similar slopes at 3 h, but at 6 h SE-sGAL animals showed a shallower slope, whereas the SE-sGLU dropped in blood pH with increasing levels of blood bacteria. This indicates improved disease tolerance following galactose supply. At the end of the study, the difference between slopes was less pronounced, but there was now a tendency for decreased health in the SE-sGAL group when bacterial levels increased. In contrast, animals in SE-sGLU reached a paralysis state with consistently low health across various levels of blood bacteria.

At euthanasia, all infected animals showed lower levels of negative acute phase reactants (albumin, cholesterol and alkaline phosphatase), but galactose supplementation alleviated drops in these markers (**Figure 2J–L**). Other liver injury markers, including aspartate aminotransferase and alanine aminotransferase, as well as kidney injury marker creatinine, were not different between the two groups (**Figure S2J–M**).

### Galactose supply reduces inflammation and enhances hepatic mitochondrial metabolism

Galactose can be converted to glucose in the liver and subsequently stored as glycogen or released directly into the bloodstream^24^. Therefore, we analyzed the hepatic transcriptome to explore metabolic and inflammatory pathways following galactose supply. The transcriptomic profiles of infected animals from uninfected ones were more distinct than that between the two infected groups (**Figure S3A**). Via gene set enrichment analysis (GSEA), SE-sGAL animals exhibited upregulation of various metabolic pathways, including OxPhos, TCA cycle, glycolysis/gluconeogenesis, as well as downregulation of various innate and adaptive immune pathways, including Th1/Th2, Th17, and IL-17 signaling (**Figure S3B and Table S2A**). Analysis of differentially expressed genes (DEGs) showed similar patterns (**Figure 2M**). The same metabolic effects of galactose were present in non-infected animals as well (**Figure S3C and Table S2A**). Taken together, the hepatic transcriptomic data further supports our conclusions from clinical parameters that galactose supply improved mitochondrial energy metabolism leading to the prevention of excessive inflammation and sepsis.

### Combined glucose restriction and glucogenic amino acid supplementation prevent sepsis and organ dysfunction

Replacing glucose with galactose enabled piglets to maintain stable blood sugar levels. However, it appeared to delay, not prevent clinical deterioration, possibly as galactose still enters glycolysis to fuel ATP production and inflammation. We therefore explored an alternative nutritional strategy to boost TCA cycle activity (to reduce inflammation) and gluconeogenesis (to ensure normoglycemia) during periods of glucose restriction. First, we screened the immune responses of *S. epidermidis*-infected THP-1 cells following supplementation of either glucose or individual or a mixture of four strictly glucogenic amino acids (GAAs): glutamate, valine, aspartate and asparagine. These GAAs were selected based on their easy conversion into TCA cycle metabolites prior to gluconeogenesis, without directing to pyruvate and lactate production. Valine and high dose of the mixture (8mM) reduced TNF-α production (**Figure 3A**), with no impact on IL-10 (**Figure S4A**) and minimal effects on gene expression (**Figure S4B**). However, all individual GAAs, as well as low (2mM) and high doses of the mixture increased ATP production after 4 hours, compared to the infected controls (**Figure 3B**). The high dose mixture increased ATP production more effectively than the addition of glucose treatment. Among the cell treatments after 6 hours, only the high dose mixture showed increased ATP production (relative to the infected controls), and with higher levels than glucose treatment (**Figure 3B**). This suggests that GAA supplementation may enhanced TCA and Oxphos during infection to produce more energy but maintained low levels of inflammation.

**Figure 3.**
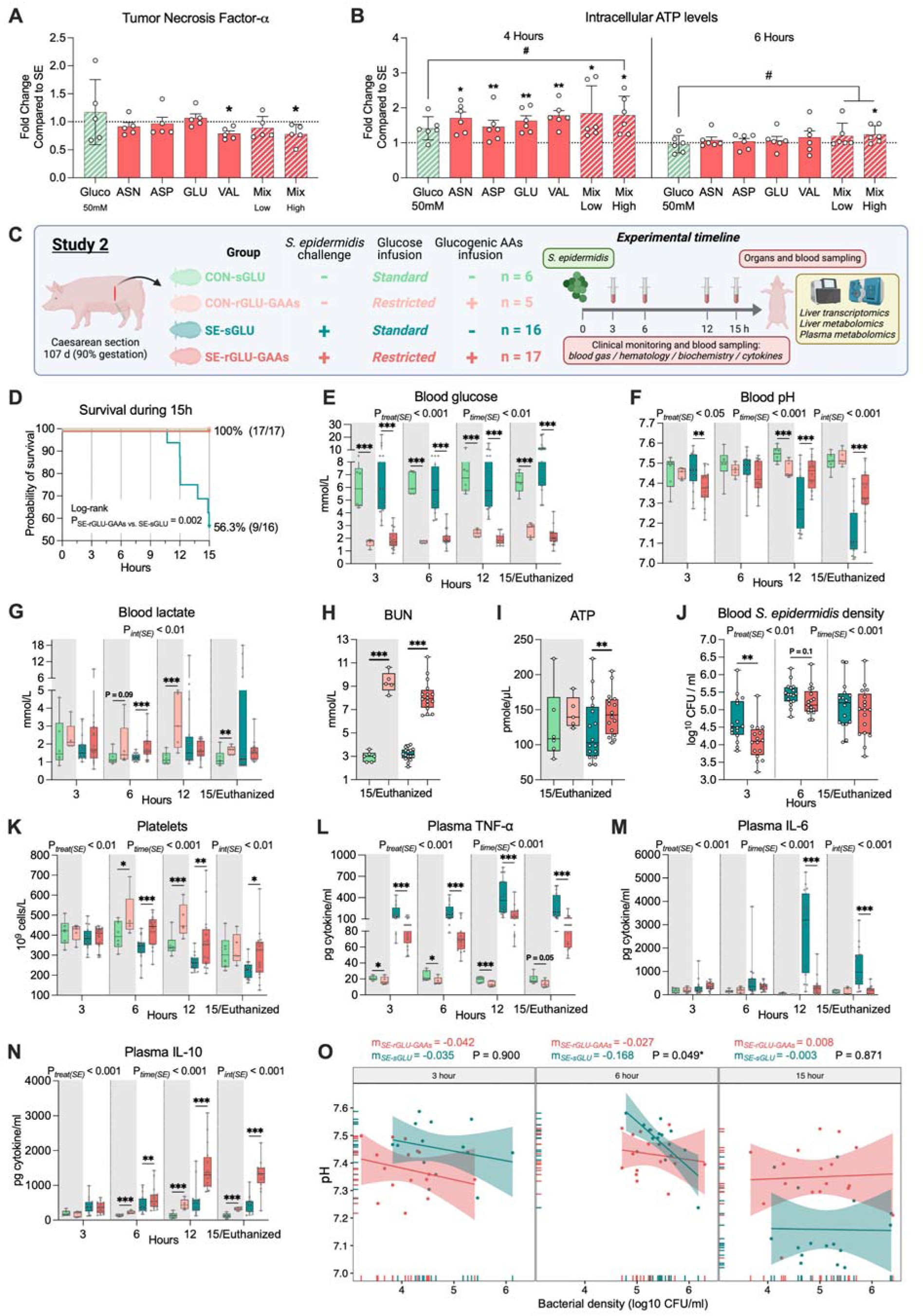
Combined glucose restriction and glucogenic amino acids supply prevent lethal sepsis and organ dysfunctions. **(A–B)** *In vitro* human macrophage-like THP1 cells exposed to live *S. epidermidis*. Levels of tumor necrosis factor-α and intracellular ATP are shown as relative fold changes in relation to *S. epidermidis* positive control. **(C)** Experimental design. Forty-four cesarean-delivered preterm piglets were randomly assigned to be exclusively nourished with parenteral nutrition containing either a combination of 1.4 % glucose (2 g/kg/d) and four glucogenic amino acids (aspartate, glutamate, asparagine, valine; each 0.5 g/kg/d) (rGLU-GAAs) or 10% glucose (sGLU; 14.4 g/kg/d). Piglets were infused with *S. epidermidis* (SE) or control saline (CON). This setup allowed for the division of piglets into four groups: CON-sGLU (n = 6), CON-rGLU-GAAs (n = 5), SE-sGLU (n = 16), and SE-rGLU-GAAs (n = 17). **(D)** Survival curves of the animals throughout the experiment, displayed as the time until humane endpoints or scheduled euthanasia. **(E–G)** Blood glucose, pH and lactate at 3, 6, 12, and 15 h post-inoculation or at euthanasia. **(H,I)** Serum blood urea nitrogen (BUN) and plasma ATP at 15 h post-inoculation or at euthanasia. **(J)** The density of *S. epidermidis* in the blood of infected piglets, determined by (CFU) assays. **(K–N)** Blood platelets, plasma TNF-α, IL-6, and IL-10) at 3, 6, 12, and 15 h post-bacterial inoculation or at euthanasia. **(O)** Reaction norm analysis was performed by plotting blood pH (as a general health indicator) against pathogen burden at 3, 6, and 15 h post-infection by linear regression. Extra sum-of-squares F Test was used to compare slopes. **Statistics: (D)** A log-rank test was used to compare survival between the SE-rGLU-GAAs and SE-sGLU groups. **(E–N)** Data at each time point were analyzed using a linear mixed-effects model, incorporating group, gender, and birth weight as fixed factors and litter as a random factor. *P-value < 0.05, **P-value < 0.01, and ***P-value < 0.001, compared between SE-rGLU-GAAs and SE-sGLU groups at the same time point. **(E–F & J–N)** Another linear mixed-effects model was employed to probe further disparities spanning the entire experimental duration, incorporating group, time, their interaction, gender, and birth weight as fixed factors, with litter and pig ID as random factors. P_treat(SE)_, P_time(SE)_, and P_int(SE)_ denote probability values for group effect (SE-rGLU-GAAs and SE-sGLU) over time, time effects, and the interaction effects between time and group in the linear mixed effects interaction model, respectively. Uninfected animals (CON) served as a reference group and were not compared directly with infected animals (SE). Statistical significance was defined as P-value < 0.05. All data were presented as box and whisker plots showing the range from minimum to maximum values. Panel C was created using Biorender.com.

Based on this, we hypothesized that a nutritional strategy of coupling glucose restriction with the supplementation of GAAs could confer better glycolysis inhibition, improve TCA cycle activity and gluconeogenesis while leading to improved infection outcomes. The same neonatal sepsis model was used with animals nourished either with standard glucose (sGLU) or restricted glucose with GAA supplementation (each amino acid 0.5 g/kg/day, rGLU-GAA, **Figure 3C**). Strikingly, 100% of SE-rGLU-GAA animals were alive at 15h post inoculation as opposed to 56% of SE-sGLU succumbing to fatal sepsis (**Figure 3D**). Further, SE-rGLU-GAAs animals maintained stable blood glucose levels of around 2 mM, in contrast to most SE-sGLU being hyperglycemic (**Figure 3E**). Clinical parameters also indicated gradual deterioration with respiratory acidosis in SE-sGLU animals from 12 h until the end of the study, indicated by severe drops of pH (**Figure 3F**), spO2, base excess, and hemoglobin and elevation of pCO2 (**Figure S4C-F**). In contrast, rGLU-GAA intervention prevented all these disturbances. Similar to galactose-nourished animals, rGLU-GAAs animals exhibited a slight, yet significant and transient, rise in lactate levels at the early phase of infection (6 h, **Figure** 3**G**). This increase suggests an enhanced conversion of the administered GAAs into pyruvate, which is then metabolized into lactate, likely through the TCA cycle. Moreover, the SE-rGLU-GAAs group at the end of the study exhibited increased blood urea nitrogen (BUN) and ATP production, suggesting the GAAs were metabolized through the TCA cycle and supported OxPhos (**Figure 3H-I**). Biochemical parameters measured at euthanasia also indicate that glucose restriction in combination with GAAs protected against injuries to the liver (lower AST and ALT) and kidneys (lower creatinine), as well as prevented impairment in hepatic synthesis of negative acute-phase reactants (higher albumin and cholesterol) (**Figure S4G–K**).

### Glucose restriction plus GAAs supply enhances both disease resistance and tolerance

With clear clinical benefits of the rGLU-GAA intervention during infection, we further examined bacterial clearance, immune, and inflammatory status to explore the associated host defense strategies. Blood bacterial loads in SE-rGLU-GAA animals were reduced at both 3 and 6 h, indicating an improved disease resistance eliminating pathogens during the early phase of infection (**Figure 3J**). This was associated with better preservation of blood leukocyte subsets (**Figure S4L–O)** and prevention of thrombocytopenia by the intervention (**Figure 3K**). Notably, different from galactose intervention, rGLU-GAAs reduced levels of TNF-α and IL-6 and elevated levels of anti-inflammatory cytokine IL-10 (**Figure 3L-N**) across all study timepoints. The differences in disease resistance and inflammatory status likely contribute to the substantial protective effects of rGLU-GAA treatment against organ injuries, which was not observed during the galactose intervention.

To evaluate disease tolerance, we again employed reaction norm analysis. At 3 h, both infected groups showed similar slopes (**Figure 3O, left**). However, by 6 h, SE-rGLU-GAAs animals had improved disease tolerance as shown with a shallow slope, in contrast to the infected controls with a steep decrease in health when bacterial burdens increased (**Figure 3O, middle**). At euthanasia, although the slopes of the reaction norms were similar between the two groups (**Figure 3O, right**), the low blood pH suggests that infected controls were in a state of immunoparalysis, whereas SE-rGLU-GAAs animals still maintained vigor. Collectively, the combined glucose restriction and GAA supplementation boosted disease resistance in early phase of infection and improved disease tolerance during the whole study period.

### rGLU-GAAs suppresses hepatic glycolysis and elevates gluconeogenesis and TCA activity

We hypothesized that protection against sepsis in SE-rGLU-GAA animals is mediated by reduced glycolysis due to glucose restriction and increased TCA cycle activity and gluconeogenesis due to the supplementation of glucogenic amino acids. First, we explored these effects via hepatic transcriptomics. The gene expression profiles indicated major impacts of both intervention and infection (**Figure S5A**). GSEA analysis between the two infected groups revealed that most metabolic pathways were upregulated in SE-rGLU-GAAs animals, including OxPhos, TCA cycle, glycolysis/gluconeogenesis, and metabolism of amino acids and fatty acids (**Figure 4A and Table S3A**). The same metabolic effects of rGLU-GAA were present in non-infected animals as well (**Figure S5B and Table S3B**). In contrast, multiple inflammation and adaptive immune pathways were suppressed. DEGs analysis revealed 3518 up-regulated and 3721 down-regulated DEGs in SE-rGLU-GAAs animals. Importantly, within 96 genes of glycolysis/gluconeogenesis, 54 genes were regulated, and most genes strictly involved in gluconeogenesis (14/16) were upregulated, whereas most genes strictly involved in glycolysis (26/35) were downregulated in SE-rGLU-GAAs animals (**Figure 4B**). This indicates that glycolysis was inhibited by glucose restriction while GAA supply markedly enhanced gluconeogenesis. Additionally, other DEG data related to TCA cycle, fatty acid β-oxidation, ketogenesis, OxPhos and amino acid metabolism as well as inflammation also support GSEA findings (**Figure S5C–F, S6A-H**).

**Figure 4.**
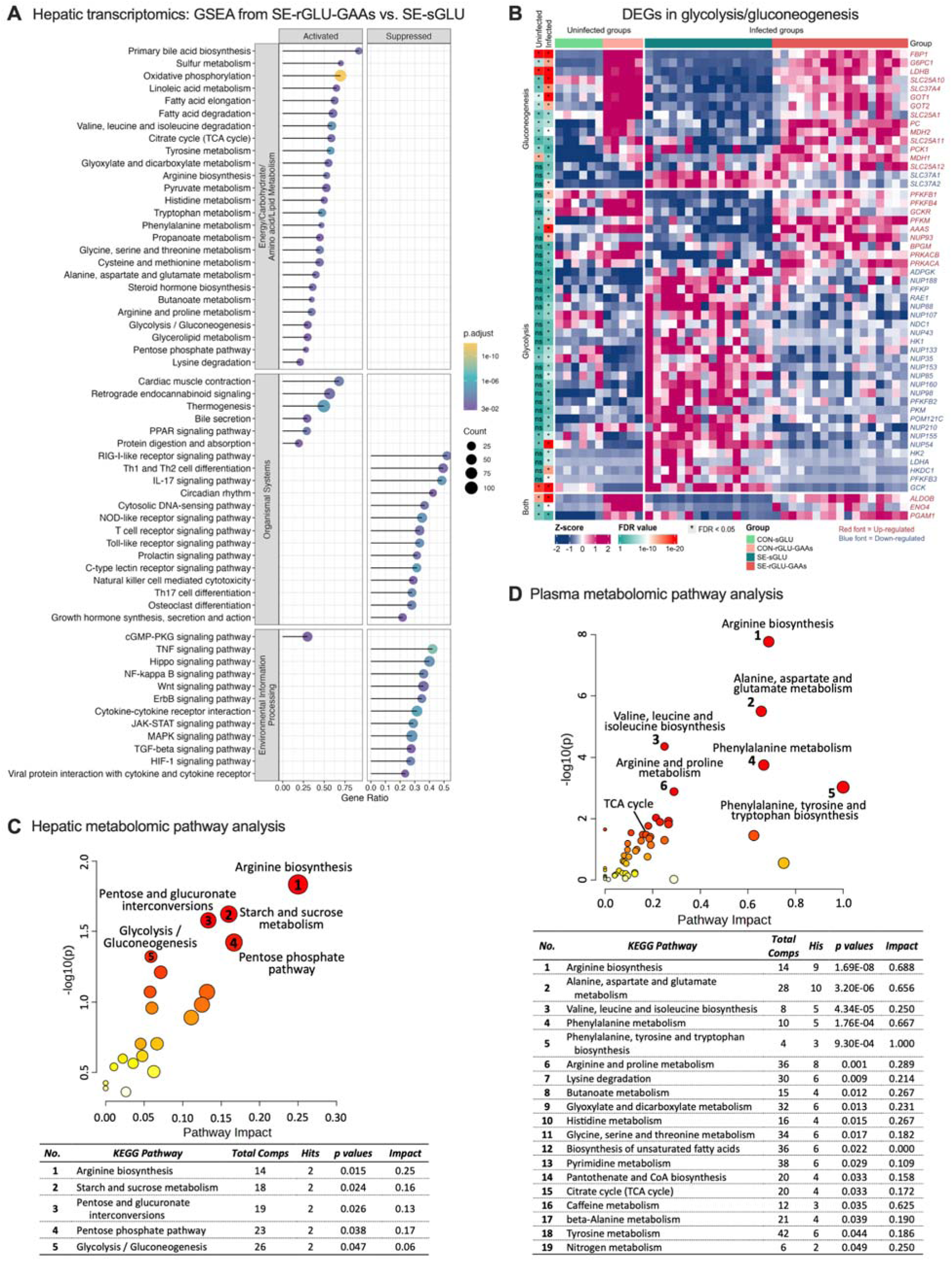
Hepatic transcriptomics and metabolomics revealed sepsis preventive effects of rGLU-GAAs supply associated with suppression of glycolysis and inflammation and elevation of gluconeogenesis. **(A)** GSEA was performed between the SE-rGLU-GAAs and SE-sGLU groups using the *Sus scrofa* (pig) KEGG knowledgebases. Significant pathways in specific categories such as energy metabolism, carbohydrate metabolism, amino acid metabolism, lipid metabolism, signal transduction, and immune system have been selected for visualization. The complete list of enriched pathways can be found in **Table S2A**. The size and color of the dots indicate the gene ratio and FDR values, respectively. **(B)** Heatmap illustrating 54 DEGs involved in the glycolysis/gluconeogenesis pathways between SE-rGLU-GAAs and SE-sGLU groups. **(C, D)** MDAs-based pathways results in hepatic tissue and plasma between SE-rGLU-GAAs and SE-sGLU groups. The color of a circle indicates the level of enrichment significance (y-axis), with red for high and yellow for low, and the size of a circle is proportional to the pathway’s impact value (x-axis). The table presents the significant pathways.

Next, we sought to confirm metabolic regulations by hepatic and plasma metabolomics. The overall hepatic metabolome with 1014 identified metabolites in 4 experimental groups was clustered together in PCA analysis (**Figure S7A**). Between the two infected groups, 77 metabolites with differential abundance (MDAs) were detected (**Table S3C**), and were associated with arginine biosynthesis and carbohydrate metabolism, including glycolysis/gluconeogenesis (**Figure 4C and Table S3D**). Specifically, carbohydrate-derived metabolites and lactic acid were reduced by SE-rGLU-GAAs, while most of the fatty acids, including carnitines and arachidonic acid dervivates were up-regulated (**Figure S7B**). These support the finding of attenuated glycolysis and increased TCA and OxPhos activities from transcriptomic data. Plasma metabolome was also explored as significant differences between hepatic and plasma metabolome during sepsis have previously reported (Khaliq et al., 2020; Oh et al., 2022). Though this analysis showed a clear clustering between the two infected groups with numerous MDAs (122 increase and 160 decrease by the intervention, **Figure S7C** and **Table S3E**), MDA-based pathway analysis revealed similar general pattern of metabolic regulations to hepatic transcriptomics and metabolomics (**Figure 4D and Table S3F, Figure S7D-F**). These included elevation of 2-oxoglutaric acid, the metabolite catabolized together with GAAs prior to entering the TCA cycle, as well as elevation of multiple glucogenic and ketogenic amino acids, ketone bodies, carnitine, and fatty acid derivatives. Collectively, data from hepatic transcriptomics and hepatic and plasma metabolomics are in agreement with each other and support findings from clinical parameters. The sepsis preventive effects of glucose restriction and glucogenic amino acid supplementation were strongly tied to a rewiring of the host energy metabolism, resulting in reduced glycolysis and enhanced TCA cycle, gluconeogenesis, amino acids metabolism, ketogenesis, and fatty acid β-oxidation (**Figure 5 and Figure S8**).

**Figure 5.**
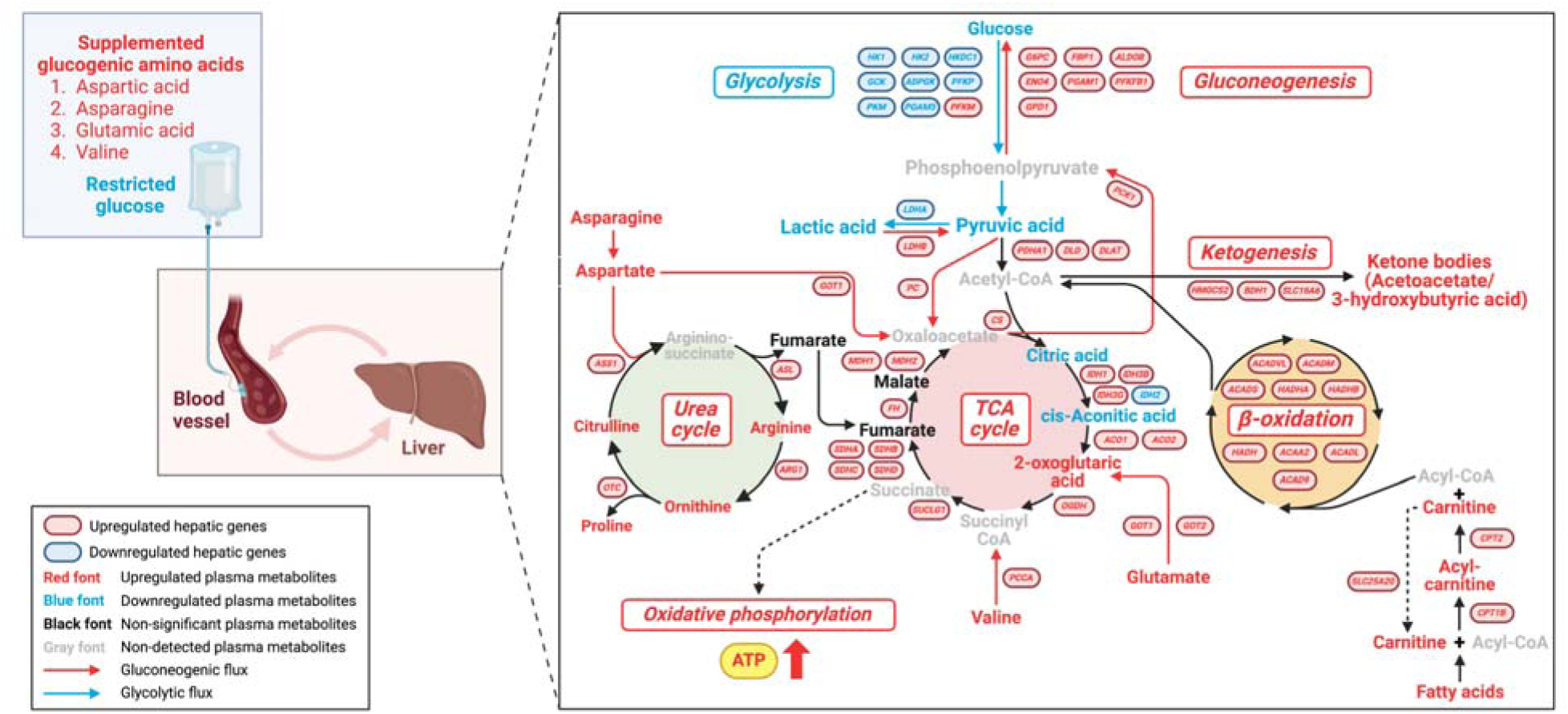
Effects of rGLU-GAAs supply on plasma metabolome during neonatal infection. Schematic representation of key metabolic alterations in cellular energy pathways (glycolysis, gluconeogenesis, TCA cycle, fatty acid β-oxidation, ketogenesis, and urea cycle) derived from integrated blood parameters, liver transcriptome profiles, and plasma metabolome data. Pathway visualization is based on genes and metabolites from KEGG knowledgebases, supplemented with additional insights from Reactome knowledgebases and the existing literature.

### GAA supply during glucose restriction elevates blood glucose and enhances both disease resistance and tolerance

Testing rGLU-GAA intervention did not separate the effect of GAAs from glucose restriction. Here, we examined the impacts of GAAs on a glucose-restricted background on glycemia and host defense (**Figure 6A**). Glucose-restricted animals were severely hypoglycemic (mean blood glucose < 1 mM) with 2/7 animals succumbed to fatal sepsis, while the GAA-supplemented animals had stable blood glucose levels around 2 mM during the 15 h of the study (**Figure 6B-C**), indicating induction of gluconeogenesis by GAAs. GAAs also prevented respiratory acidosis (higher blood pH, lower pCO2, and unaltered lactate, **Figure 6D–F**), while increased BUN levels suggest increased amino acid catabolism (**Figure 6G**). Importantly, GAAs enhanced bacterial clearance at both 3 and 6 h, as well as better blood leukocyte replenishment and reduced inflammation at the later phase of the experiment (**Figure 6 H–L**). Subsequent reaction norm analysis between the two groups showed similar slopes at 3-6 h, but GAA animals again had improved disease tolerance at the end of the study (**Figure 6M)**. Together, this experiment confirmed that GAAs directly contributed to the clinical benefits of the rGLU-GAA intervention, most likely via enhancement of both disease resistance and tolerance, as well as improved hepatic gluconeogenesis. Importantly, while our earlier research established a link between hyperglycemia and heightened sepsis severity,^6,7^ the current findings indicate that avoidance of severe hypoglycemia also contributed to sepsis prevention. The higher glucose levels in SE-rGLU-GAAs vs. SE-rGLU animals were not exogenous but stemmed from gluconeogenesis induced by supplying GAA during glucose restriction.

**Figure 6.**
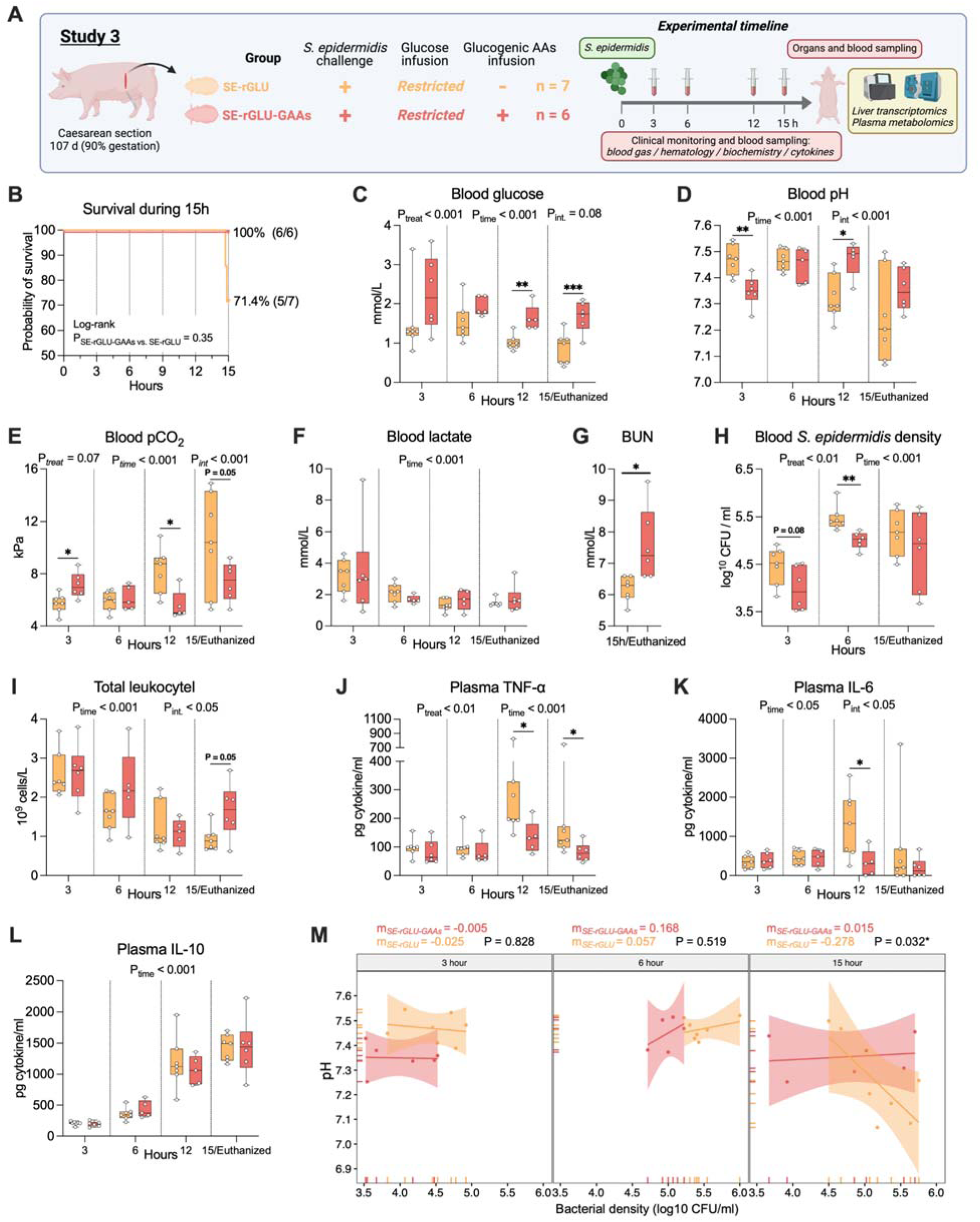
Glucogenic amino acid supplementation is sufficient to modulate clinical parameters and induce metabolic rewiring during sepsis. **(A)** Schematic overview of the experimental design. Thirteen preterm piglets delivered by cesarean section were randomly assigned to groups exclusively nourished with parenteral nutrition containing restricted glucose levels (1.4 %, 2 mg/kg/day), without (rGLU, n=7) vs. with supplementation of four glucogenic amino acids (aspartate, glutamate, asparagine, and valine, each 0.5 g/kg/day, rGLU-GAAs, n=6). All piglets were infused with *S. epidermidis* (SE). **(B)** Survival curves of the animals throughout the experiment, displayed as the time until humane endpoints or scheduled euthanasia. **(C–F)** Blood gas parameters (blood glucose, pH, pCO_2_, and lactate) at 3, 6, 12, and 15 h post-inoculation or at euthanasia. **(G)** Blood urea nitrogen (BUN) at 15 h post-inoculation or at euthanasia. **(H)** Blood *S. epidermidis* density, determined by colony-forming units (CFU) assays. **(I–L)** Blood immune parameters (total leukocytes, plasma TNF-α, IL-6, and IL-10) at 3, 6, 12, and 15 h post-inoculation or at euthanasia. **(M)** Reaction norm analysis, performed using blood pH (as a general health indicator) at 3, 6, and 15 h and plotted against the pathogen burdens at the same time point by linear regression. Extra sum-of-squares F Test was used to compare slopes. **Statistics: (C–L)** Data at each time point were analyzed using a linear model, incorporating group, gender, and birth weight as fixed factors. *P-value < 0.05, **P-value < 0.01, and ***P-value < 0.001, compared between SE-rGLU-GAAs and SE-rGLU groups at the same time point. **(C–F & H–L)** A linear mixed-effects model was employed to probe further disparities spanning the entire experimental duration, incorporating group, time, and their interaction as fixed factors, with pig ID as random factors. P_treat_, P_time_, and P_int_ denote probability values for group effect (SE-rGLU-GAAs vs. SE-rGLU) over time, time effects, and the interaction effects between time and group in the linear mixed effects interaction model, respectively. Uninfected animals (CON) served as a reference and were not included in the statistics. Statistical significance was defined as P-value < 0.05. All data were presented as box and whisker plots showing the range from minimum to maximum values. Panel A was created using Biorender.com.

### GAA supply during glucose restriction elevates gluconeogenesis

Next, we also sought to confirm these GAA effects via hepatic transcriptomics and plasma metabolomics. Distinct hepatic transcriptome profile between the two groups was shown by both PCA (**Figure 7A**) and DEG analysis with 189 upregulated and 136 downregulated genes in SE-rGLU-GAAs group. From both GSEA and DEG enrichment analyses, GAAs upregulated OxPhos, TCA cycle, glycolysis/gluconeogenesis, amino acid and lipid metabolic pathways, while dampened several inflammatory pathways (**Figure 7B, S9A-B and Table S**4**A**). For plasma metabolomics, PCA also revealed distinct clustering patterns between the two groups (**Figure 7C**), while MDA analysis showed increased levels of most amino acids, acylcarnitines, ketone body, keto acid and fatty acids in SE-rGLU-GAAs animals (**Figure 7D and Table S4B**). It was clear that SE-rGLU-GAAs vs. SE-sGLU comparison (study 2) had much more profound changes in hepatic transcriptome and plasma metabolome than SE-rGLU-GAAs vs. SE-rGLU comparison (study 3). This implies the metabolic rewiring impacts were mainly mediated by glucose restriction. Regardless, the additive effects of GAAs in the background of glucose restriction on disease resistance and tolerance, as well as gluconeogenesis, TCA cycle and systemic inflammation appeared to be critical to both sepsis prevention and glucose homeostasis.

**Figure 7.**
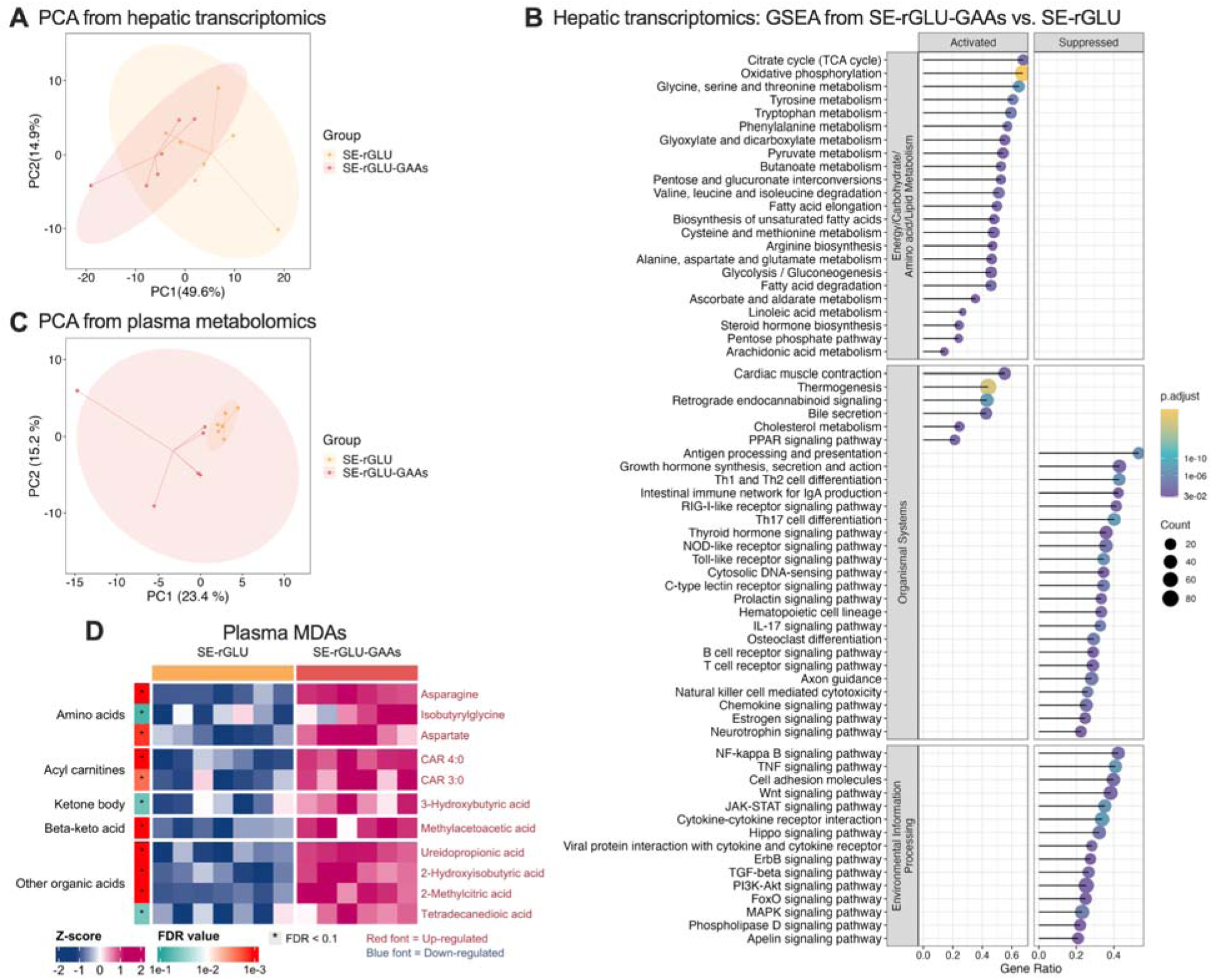
GAA supply during glucose restriction enhances gluconeogenesis and reduces inflammation during neonatal infection. **(A,C)** Hepatic Transcriptomics and plasma metabolomics PCA scores plots of the first two principal components. **(B)** GSEA, performed between the SE-rGLU-GAAs and SE-rGLU groups using the *Sus scrofa* (pig) KEGG knowledgebase. Significant pathways in specific categories such as energy metabolism, carbohydrate metabolism, amino acid metabolism, lipid metabolism, signal transduction, and immune system have been chosen for presentation. The complete list of enriched pathways can be found in **Table S2**. The size and color of the dots indicate the gene ratio and FDR values, respectively. **(C)** Heatmap illustrating DEGs involved in the energy metabolism-related pathways between SE-rGLU-GAAs and SE-rGLU groups. **(D)** Heatmap illustrating DEGs involved in the immune/inflammation-related pathways between SE-rGLU-GAAs and SE-rGLU groups.

## Discussion

Combining human birth cohort data, *in vitro* screening and *in vivo* intervention studies with a newborn piglet infection model, we consistently documented a tight connection between energy metabolism and host defense in both healthy term human newborns and preterm piglets with high infection risks. Data from the COPSAC_2010_ mother-child cohort indicate that normal variation in TCA cycle activity was associated with altered risk of common bacterial infections in childhood (otitis media, tonsilitis and cold symptoms), systemic inflammation and immune functions. These findings are in alignment with previous literature and indicate that activity of the TCA cycle not only correlates with anti-inflammatory/tolerogenic immune responses, but also better infection resilience^9–11^. However, the association between energy metabolism and immunity appears to be more complex in those born prematurely or at a low birthweight. In hospitalized neonates, hypoglycemia is among the most common metabolic disturbances shortly after birth, which is partly derived from their low levels of fat and glycogen stores^27^. On the other hand, premature newborns are often insulin resistant, which causes high risks of hyperglycemia upon nourishment with glucose-rich PN, increasing their risk of infections developing into sepsis^6,7,28,29^. Hence, newborn health maintenance upon infection requires a delicate balance of host energy metabolism to control defense strategies with appropriate levels of inflammation^5–7^. Here using a neonatal sepsis model, we identified two novel nutritional strategies to boost TCA cycle activity via galactose and GAA supplementation, which enhanced defense strategies in distinct manners to promote infection survival.

Despite being a natural part of lactose in breastmilk, and released in the gut and absorbed after breastfeeding, galactose has not been used directly as a carbohydrate source for newborn infants. Immune cells cultured in galactose-containing medium showed suppressed glycolytic activity and lactate production^30^, as it must first be metabolized before entering into glycolysis, or released as glucose^31^. Studies in healthy adults have also demonstrated that intravenous galactose administration may lead to an increase in hepatic glycogen and improved glucose homeostasis. In the current study, the supply of galactose, instead of glucose enhanced disease tolerance during early phase of infection and overall anti-inflammatory response over the infection course and prevented hyperglycemia, improving survival and clinical outcomes. However, important markers of liver and kidney injuries (liver enzymes, albumin, cholesterol, creatinine) were minimally affected by this, together with limited changes in hepatic metabolic rewiring measured by transcriptomics at the end of the study. It thus appeared that substituting galactose for glucose still resulted in some downstream glycolytic activity, delaying rather than preventing the progression of excessive inflammation. In clinical settings, sepsis delaying effects of the galactose intervention may provide important window of opportunity for other adjunct therapies (e.g., antibiotics or inotropes) in infected vulnerable newborns prior to clinical deterioration.

In contrast, the alternative approach of combining glucose restriction and GAA supplementation effectively limited glycolysis, enhanced OxPhos and gluconeogenesis, and markedly improved infection survival and protection against organ damage. The metabolism of GAAs via TCA cycle can provide energy via OxPhos for the liver and immune system, affecting their inflammatory response and via hepatic gluconeogenesis help maintain a supply of glucose for other vital organs such as the brain. Preterm infants possess the required enzymes for gluconeogenic activities^32,33^ and it is possible that glucose homeostatic control during neonatal infection in preterm infants can be achieved by restricting PN glucose supply and GAA supplementation. Importantly, this intervention did not completely inhibit inflammation and the treated animals were capable of mounting sufficient disease resistance response to better eliminate invading bacteria during the first six hours of infection. Strikingly, disease tolerance mechanisms were also enhanced by the intervention at six hours and maintained until the end of the study. Clearly, infection survival was not correlated with the host capacity to eliminate invading pathogens, but rather dictated by the balance of host defense strategies. To our knowledge, this was the first study showing simultaneous enhancement of both disease tolerance and resistance during infection.

Enhanced disease tolerance exerted by glucose restriction and GAAs was also associated with improved hepatic energy metabolic pathways, not only TCA cycle and OxPhos but also fatty acid oxidation, and ketogenesis. These pathways have been shown to play critical roles in providing alternative energy for vital organs (e.g., brain and heart) and protecting against inflammation-induced tissue damage during sepsis.^34^ Further, ketone bodies, particularly beta-hydroxybutyrate can exhibit potent anti-inflammatory effects^35,36^. Finally, all these effects are strikingly similar to the regulated metabolic pathways and defense strategies we have previously demonstrated in preterm pigs that survived neonatal infection, relative to those succumbed to sepsis^5^.

In conclusion, our results not only strongly imply connections between newborn metabolic regulation and host infection defense but also suggest potential therapies that may be lifesaving for vulnerable newborns with suspected infections. Both galactose and glucose restriction plus GAA supplementation conferred protective effects during serious bloodstream infections and modulated newborn defense strategies. Given the obvious safety of galactose, phase 1 trials in vulnerable newborns requiring monosaccharide infusion may be best way to determine if human newborns are capable of utilizing galactose effectively, either alone or in some combination with glucose. Our encouraging results may therefore directly pave the way for clinical trials testing the feasibility and efficacy of these interventions.

### Limitations of the study

With the human cohort data, we were only able to establish associations between semiquantitative levels of plasma metabolites and infection burden, that do not prove causal links between energy metabolism and infection susceptibility. Further, the granular nature of the data allowed the distinction between suspected bacterial and viral infections although it must be noted that the cohort comprised healthy children and thus had a very low rate of serious infections requiring hospitalization. Likewise, although our animal experiments showed the clinical benefits of novel nutritional strategies, certain limitations must be acknowledged. First, our preterm sepsis model with *S. epidermidis* mimics neonatal nosocomial coagulase-negative staphylococci infection in preterm infants, and therefore, it is possible that different bacterial species could elicit various defense strategies. Second, due to resource constraints and the limited volume of blood collected during the study, a more detailed characterization of host defense, immune, and metabolic responses at various time points during infection was not possible, limiting a comprehensive understanding of host metabolic changes over time. Finally, our latter experiments focused on the effects of four selected strictly glucogenic amino acids. Therefore, the mechanistic impacts of each amino acid in this setting are unknown. It is important in future research to explore optimal dosage and the contributing effects of each amino acid during neonatal infection.

## METHODS

### Key resources table

| REAGENT or RESOURCE | SOURCE | IDENTIFIER |
| --- | --- | --- |
| **Bacterial strain** |  |  |
| *Staphylococcus epidermidis* | Isolated from a sepsis infant | WT-1457 |
| **Chemicals and reagents** |  |  |
| D-(+)-Galactose | Sigma | G5388 |
| L-Aspartate | Sigma | A7219 |
| L-Asparagine | Sigma | A4159 |
| L-Valine | Sigma | V0513 |
| L-Glutamate | Sigma | G8415 |
| Doxapram | Carinopharm |  |
| Flumazenil |  |  |
| Zoletil 50 Vet. | Virbac | 568527 |
| Xylazine |  |  |
| Ketamine |  |  |
| Pentobarbital | Alfasan | 088672 |
| Kabiven | Fresenius Kabi | 052505 |
| Heparin | Panpharma | ETISM788-1 |
| **Critical Commercial Assays** |  |  |
| ATP Assay Kit | Sigma-Aldrich | MAK190 |
| Porcine TNF-α DuoSet ELISA Kit | R&D | DY690B |
| Porcine IL-6 DuoSet ELISA Kit | R&D | DY686 |
| Porcine IL-10 DuoSet ELISA Kit | R&D | DY693B |
| RNeasy mini kit | Qiagen | 74106 |
| Human TNF alpha Uncoated ELISA Kit with Plates | ThermoFisher | 88-7346-22 |
| **Experimental Models: Organisms/Strains** |  |  |
| Preterm piglets | crossbred, Landrace × Yorkshire × Duroc | |
| **Deposited data** |  |  |
| RNA sequencing data | NCBI | GEO: GSEXXXXX |
| **Software and Algorithms** |  |  |
| GraphPad Prism 9 | GraphPad | https://www.graphpad.com/scientific-software/prism/ |
| R version 4.2.3 | R project |  |
| MetaboAnalystR 6.0 | Metaboanalyst | https://www.metaboanalyst.ca/ |
| **Others** |  |  |
| GEM Premier 3000 | Instrumentation Laboratory | GEM Premier 3000 |
| ADVIA 2120i Hematology System | Siemens | ADVIA 2120i System |
| ADVIA 1800 Chemistry System | Siemens | ADVIA 1800 System |

### Resource availability

#### Lead contact

Further information and requests for resources and reagents should be directed to and will be fulfilled by the lead contact. Dr. Duc Ninh Nguyen.

#### Materials availability

This study did not generate new unique reagents.

#### Data and code availability

The raw and processed RNA-seq data generated from this study was deposited in the NIH Gene Expression Omnibus, with accession number GSE263512. The original and normalized hepatic and plasma metabolomics data were attached to the supplementary data. The pipeline for the analysis is available on GitHub (https://github.com/Pharmaco-OmicsLab/GAA_Sepsis). Any additional information required to reanalyze the data reported in this paper is available from the lead contact upon request.

## HUMAN DATA

### Study Population

The Danish population-based COPSAC_2010_ mother–child cohort has been comprehensively described in earlier publications, including detailing participant baseline characteristics and the enrollment process^16,17^. Briefly, 736 pregnant women were recruited at 24 weeks of gestation, and their children were monitored longitudinally at the COPSAC research clinic until the age of 3 years, with extended follow-up visits at 4, 5, 6, 8 10 and 13 years. This included 9 planned visits at age 0-3 years as well as acute care visits, with a primary focus on common infections and atopic conditions. The pregnant women participated in a double-blind randomized controlled trial using a 2×2 factorial design, receiving either fish oil capsules containing 2.4 g of n-3 long-chain polyunsaturated fatty acids or a placebo (Clinicaltrails.gov: NCT00798226). Additionally, 623 participants were enrolled in another RCT comparing high-dose vitamin D supplementation (2800 IU/day) with a standard dose (400 IU/day) (Clinicaltrails.gov: NCT00856947). The study received approval from the Danish Ethics Committee (H-B-2008-093) and the Danish Data Protection Agency (2008-41-2599).

Infections were recorded through acute visits and daily diaries during the first 3 years and included pneumonia, tonsillitis, acute otitis media, gastroenteritis, cold symptoms and fever. Diagnoses were confirmed by COPSAC physicians at scheduled and acute care visits based on parental interviews, medical records and registry data^37^.

### Metabolomics Profiling

Metabolite data were obtained from plasma samples collected from the mother at time of birth and from the child at 6 and 18 months of age. LC-MS metabolite characterization was performed by Metabolon, Inc. (Durham, NC, USA). The analytical method used for these samples has been described ^38,39^.The identification level follows the criteria described previously^40^. A data normalization step was applied to correct variations due to inter-day instrument tuning differences by aligning median values to one and proportionally normalizing each data point. For samples involving multi-day sample runs, batch normalization was performed to adjust for minor technical variations between batches. All data were transformed using the natural logarithm before statistical analysis.

### Immune Profiling

At 18 months of age blood was collected in heparinized tubes and analyzed within 4 hours. The details of the blood collection and immune characterization has been published elsewhere in detail^15^. *Brefly, for flow Cytometry* cell subsets were defined based on specific marker combinations and side scatter (SSC) properties as follows: granulocytes (SSC^int/high^), neutrophils (CD16^+^ SSC^int/high^), eosinophils (CD16^-^ SSC^high^), B cells (CD19^+^), T cells (CD3^+^), CD4 T cells (CD3^+^CD4^+^), CD8 T cells (CD3^+^CD8^+^), activated CD4 T cells (CD3^+^CD4^+^CD25^+^CD127^high/+^), activated CD8 T cells (CD3^+^CD8^+^CD25^+^CD127^high/+^), Tregs (CD3^+^CD4^+^CD25^+^CD127^low/-^), γδT cells (CD3^+^γδTCR^+^), invariant NKT cells (iNKT, CD3^+^Vβ24Jα18^+^), classical monocytes (CD3^-^ CD19^-^CD14^+/high^CD16^-^), intermediate monocytes (CD3^-^CD19^-^CD14^+/high^CD16^+^), inflammatory monocytes (CD3^-^CD19^-^CD14^int/-^CD16^+^), BDCA-1 dendritic cells (CD3^-^CD19^-^CD14^-^CD16^-^ BDCA-1^+^), BDCA-3 dendritic cells (CD3^-^CD19^-^CD14^-^CD16^+^CD1c^-^BDCA-3^high^), plasmacytoid dendritic cells (CD3^-^CD19^-^CD14^-^BDCA-2^+^), NK cells (CD3^-^CD56^+^), CD56^dim^ NK cells (CD3^-^ CD56^dim^), and CD56^bright^ NK cells (CD3^-^CD56^bright^). Total counts of leukocytes, monocytes, dendritic cells, innate immune cells, and adaptive immune cells were also enumerated. Data were collected on a BD FACSCanto II flow cytometer and analyzed using FlowJo v7.6.5 (TreeStar) with a predefined gating strategy^15^.

*For Stimulation Assays,* whole blood was diluted 10-fold in media and stimulated in parallel with a range of pre-titrated stimuli^15^. Following 24 hours of incubation at 37°C and 5% CO, supernatants were collected and stored at −80°C. Stimulation agents included: *E. coli* derived lipopolysaccharide (LPS, InvivoGen) (500 ng/mL), synthetic dsRNA analog Poly(I:C) (100 µg/mL), imidazoquinoline (R848, 2 µg/mL) and peptidoglycan (PGN, 25 µg/mL, all InvivoGen). 1-Hydroxy-2-methyl-2-buten-4-yl 4-diphosphate (HDMAPP, 1 µg/mL, Echelon Biosciences) and Staphylococcal enterotoxin B (SEB, 200 ng/mL, Sigma-Aldrich). Cytokine and chemokine concentrations were quantified using MesoScale Discovery (MSD) pre-coated plates, including Human IFN-beta Tissue Culture Kit, Human TGF-β1 Kit, Cytokine Panel 1 V-PLEX, Prototype Human 3-plex, and MULTI-SPOT 10 Spot Special Order Human 9-plex. Undetectable levels were assigned values at half the minimum detectable concentration for each analyte. Cytokine stimulation ratios were calculated by dividing the stimulated value to the unstimulated and all values were then z-score transformed before analysis.

### Statistical Analyses

Infection risk was analyzed by a quasi-poisson regression model estimating the incidence risk ratio using number of infection episodes as count data. Time to infection was calculated by a cox-proportional hazard model while the immune data were analyzed by a general linear regression model estimating beta-coefficients using the continuous cell count or cytokine levels. Analyses from birth, were adjusted for possible confounders such as the randomization in the two RCTs, sex, social circumstances, smoking during pregnancy, gestational age, birthweight, delivery mode, neonatal hospitalization, maternal antibiotic use during pregnancy, maternal asthma, birth season, maternal age, BMI, duration, and presence of siblings while at 6 and 18 months covariates were reduced to participation in RCTs, sex, social circumstances, child BMI, and any infections 14 days previously, based on directed acyclic graphs and previous similar analysis on the impact of metabolite levels at birth on infection outcomes^41^. Analyses were conducted using Stata (version 14.2), with significance defined as P < 0.05.

## IN VITRO EXPERIMENTS

The human monocyte-like THP-1 cell line was obtained from ATCC (TIB-202) and cultivated in RPMI-media (ThermoFisher 72400047) with 10% FCS (ThermoFisher A5670502). Prior to stimulation, cells were seeded in 24 well plates, 2.5 x 10^5^ cells/well and differentiated into macrophage-like cells by treatment with Phorbol-12-myristate-13-acetate (Sigma, 524400, 180 nM) for three days. Following differentiation, cells were infected with *S. epidermidis*, MOI 0.5 for 4 or 6 hours, with and without treatment with glucose (10/50mM), galactose (10/50mM), asparginine (2mM), aspartate (2mM), glutamate (2mM), valine (2mM) or an amino acid mixture (2/8mM). Afterwards, intracellular ATP was determined using the BacTiter Glow ATP assay (Promega, 74106) and cytokine secretion was determined using commercial ELISA kits (ThermoFisher 88-8086-86, 88-7106-86, 88-7066-86) according to manufacturer’s instructions. Total RNA was isolated using the RNeasy kit according to manufacturer’s instructions (Qiagen 7404). cDNA was synthesized from 2 µg of RNA using a High-Capacity cDNA Reverse Transcription Kit (Thermo Fisher, Waltham, MA, USA). Real-time quantitative PCR (RT–qPCR) was performed with 10-fold diluted cDNA using the LightCycler 480 SYBR Green I Master kit on a LightCycler 480 (both Roche, Basel, Switzerland). Samples were analysed in duplicate using cytokine-specific primers (Supplementary Table S5). Gene expression was normalised to the expression of beta-actin using the 2-△△CT method. Data was normalized to *S. epidermidis* infected cells cultured in uncomplemented media and all statistical comparisons between treatments were done by either paired T-test (for normally distributed data) or Wilcoxon’s signed rank test. Analyses were conducted using Stata (version 14.2), with significance defined as P < 0.05.

## EXPERIMENTAL ANIMAL STUDIES

### Study approval

The Danish Animal Experiments Inspectorate approved all animal studies and experimental procedures under license number 2020-15-0201-00520. These approvals are in accordance with EU Directive 2010/63, which governs the legislation for the use of animals in research.

### Animals

Seventy-four crossbred preterm piglets (Landrace x Large White x Duroc) were delivered by cesarean section at day 106 (90% gestation, term at day 117) from three healthy pregnant sows (21, 23, and 27 piglets per sow). The anesthesia and surgical procedures of sows are described in detail elsewhere^23^. After delivery, the newborn piglets were immediately transferred to preheated (37°C) newborn incubators with a supplementary oxygen supply (1–2 L/min). When needed, animals were resuscitated by Doxapram and Flumazenil (0.1 ml/kg for each, intramuscular injection), tactile stimulation, and manual noninvasive positive pressure ventilation, applied as necessary until achieving respiratory stability. Furthermore, each piglet was equipped with a vascular catheter inserted into the dorsal aorta via the transected umbilical cord, facilitating the administration of PN, bacterial inoculation, and blood sampling. Three piglets were euthanized due to failure of jugular vein puncture (animal number: C100), catheter problem (animal number: C124), and unsuccessful resuscitation (animal number: C151) and were excluded from the study.

### *S. epidermidis* culture preparation

Originating from frozen stock, the *S. epidermidis* bacteria, isolated from a septic infant, were cultured in tryptic soy broth overnight. Following this incubation, bacterial density was accurately gauged using spectroscopic techniques. Based on these measurements, a working solution (10^9^ CFU/kg) was formulated by diluting the culture with a precise volume of sterile saline ^23^.

### Parenteral nutrition preparation

Four specialized formulations of parenteral nutrition (PN) were developed from the Kabiven infusion formula (Fresenius-Kabi). For PN with standard glucose (sGLU) provision, the glucose chamber was emptied, and a specific volume of 50% glucose solution was added into the glucose chamber to reach 10% glucose PN with infusion rate of 6 mL/kg/h, equivalent to 14.4g/kg/d glucose, similar to PN used for preterm infants.^42^ For PN with galactose supply with standard glucose equivalent level (sGAL), glucose chamber was emptied and galactose solution was added to reach 10% galactose PN (14.4 g/kg/d). For glucose-restricted (rGLU) PN, a precise volume of glucose was removed from the PN glucose chamber to reach a final glucose content of 1.4% (2 g/kg/d with an infusion rate of 6 mL/kg/h). For PN with glucose restriction and supplementation of four glucogenic amino acids (rGLU-GAAs), a precise volume of four glucogenic amino acids solution was added in the glucose-restricted PN solution to reach the infusion rate of 0.5 g/kg/d for each amino acid (aspartate, glutamate, asparagine, valine).

### Animal experimental procedures and intervention

Approximately two hours after the cesarean section, all animals were randomly stratified based on birth weight and gender into groups receiving one of the four types of PN: sGLU, sGAL, rGLU, and rGLU-GAAs. Animals were then inoculated with live *S. epidermidis* (SE) or control saline via an interatrial infusion over three minutes. During the post-inoculation period, animals reaching predefined humane endpoint (lethal sepsis criteria, including arterial blood pH of ≤ 7.1 and clinical signs of deep lethargy, discoloration, and tachypnea) were euthanized for blood sampling and tissue collection. To standardize sample collection times and improve animal welfare, we adjusted the experimental duration for the second and third litters to 15 h. This modification was prompted by initial findings where animals in the first litter received intensive care for up to 20 h post-inoculation. In contrast, all animals in the sepsis control group (SE-sGLU) were humanely euthanized before 18 h. Five out of six animals in the litter, one from the SE-sGAL group and one out of three animals from the SE-rGLU-GAAs group in the same litter, were humanely euthanized before 20 h.

Blood samples were collected at 3, 6, and 12 h post bacterial challenge and at the study end via the arterial catheter for blood gas analysis and hematology, as well as plasma collection and storage for cytokine measurements. At 3 and 6 h and at the study end, blood samples were drawn through the jugular vein puncture for bacterial enumeration. Serum and plasma samples at euthanasia were used for serum biochemical analysis and plasma metabolomic analysis.

At humane endpoint or 15 h of post-inoculation, all piglets were deeply anesthetized with a Zoletil mixture (0.1 ml/kg), which included Zoletil 50 (125 mg tiletamine, 125 mg zolazepam), xylazine (6.25 ml xylazine 20 mg/ml), ketamine (1.25 ml ketamine 100 mg/ml), and butorphanol (2.5 ml butorphanol 10 mg/ml), and were subsequently euthanized with an intracardiac injection of barbiturate. The liver was collected and preserved (frozen) during necropsy for liver transcriptomic and metabolomic analysis.

### Blood gas, hematology, inflammatory markers, serum biochemistry, and plasma ATP measurement

Blood samples at 3, 6, 12, and 15 h underwent routine blood gas analysis utilizing the GEM Premier 3000 (Instrumentation Laboratory, USA) for blood pH, pCO_2_, oxygen saturation, base excess, glucose, and lactate measurement. Hematological assessments were conducted with the ADVIA 2120i Hematology System (Siemens, Germany). For the quantification of plasma cytokines, TNF-α, IL-6, and IL-10 were analyzed using porcine-specific DuoSet enzyme-linked immunosorbent assays (R&D Systems). Serum biochemistry evaluations were executed using the ADVIA 1800 Chemistry System (Siemens, Germany). Additionally, extracellular ATP level was measured by the ATP Colorimetric Assay Kit (Sigma-Aldrich).

### Hepatic transcriptome profiling and analysis

RNeasy mini kit (QIAGEN, US) was used to extract liver RNA from all piglets. Gene expression was profiled by utilizing whole-transcriptome shotgun sequencing. Library preparation and sequencing were carried out by NOVOGENE services (Cambridge, UK), and bioinformatic pipeline was adapted from a previously published study. ^43,44^ DEG analysis was performed using the *DESeq2* package version 1.38.3.^45^ We excluded lowly expressed genes, defined as having a raw count of less than 10 less than 4, 5, and 6 samples in Experiment 1, 2 and 3, respectively. Litter was added to adjust the model. To robustly estimate the fold change (FC), we applied the *lfcShrink* function with *ashr* estimator.^46^ A false discovery rate (FDR) cut-off of 0.05 was implemented to obtain DEGs. Genes were ranked by the FC (in log scale) before pathway analyses. The gene set enrichment analysis (GSEA) module in the *clusterProfilter* package version 4.0.2^47^ was utilized with *Sus scrofa* Kyoto Encyclopedia of Genes and Genomes (KEGG) knowledgebases to elucidate pathway-level disturbances. Down- or up-regulated pathways were defined based on a normalized enrichment score (NES) with negative or positive values, respectively. The FDR cut-off of 0.05 was considered to determine significant pathways. To create a comprehensive visualization of specific genes within GSEA-enriched pathways, DEGs were identified based on the KEGG, Biological Processes aspect of Gene Ontology (GO:BP), Reactome, and Hallmark knowledgebases, with additional insights from the literature. The relative expression of DEGs was then visualized through heatmaps generated using the *ComplexHeatmap* package version 2.15.1.^48^ The analysis of liver transcriptomics data was performed in R 4.2.3.

### Hepatic and plasma metabolome profiling and analysis

Untargeted metabolomics analysis was conducted using ultra-performance liquid chromatography-mass spectrometry (UPLC-MS) through the services provided by Creative Proteomics (Shirley, NY, USA), with detailed methods described previously^5^. For hepatic metabolome, there were 5614 and 5067 features in positive and negative ion modes were detected. Among them, 1098 putativelly annotated features, assigned by the vendor and cross-examined by our team, were eventually used for subsequent analyses. For plasma metabolome, a total of 754 putatively annotated features in POS and NEG ion modes were used for downstream analyses. All raw data with annotations were described in Table S6. The low repeatability annotated features were excluded, whose relative standard deviation (RSD) was ≥ 25% in the pooled QC samples. Prior to statistical analysis, data was median-normalized and log10-transformed using *MetaboAnalystR* package version 4.0.0 in R version 4.2.3.^49^ A linear mixed-effects model was conducted, incorporating group, gender, and birth weight as fixed factors and litter as a random factor, using *lme4* packages.^50^ An FDR cut-off of 0.1 was applied to derive molecules with differential abundance (MDAs). MDAs-based pathway analysis, which integrated hypergeometric test and out-degree centrality, was performed in the MetaboAnalyst 6.0 (https://www.metaboanalyst.ca) with *Sus scrofa* KEGG knowledgebase. Pathways with a P-value cut-off of 0.05 and more than one significant hit were considered statistically significant.

### Statistics

All statistical analyses of non-omics data were executed via R version 4.3.2 unless otherwise stated. For discerning differences at specific intervals (3, 6, 12, and 15 h), continuous data underwent analysis via a linear mixed-effects model, incorporating group, gender, and birth weight as fixed factors and litter as a random factor, using *lme4* package.^50^ Another linear mixed-effects model was employed to probe further disparities spanning the entire experimental duration. This model integrated group, time, their interaction, gender, and birth weight as fixed factors, with litter and pig ID as random factors, using *lme4* package.^50^ Normal distribution, variance homogeneity of residuals, and fitted values were assessed. The data that did not conform to a normal distribution were logarithmic transformed. If transformation did not achieve approximate log-normal distribution, the non-parametric Mann-Whitney U test was used instead. To examine different strategies in the infection response, reaction norm analysis was performed using blood pH as a readout for health at 3, 6, and 15 h and plotted against the pathogen burdens at the same time point by linear regression. Extra sum-of-squares F Test was used to compare slopes. Statistical significance was defined as P-value < 0.05.

## Acknowledgments

The authors thank Thomas Thymann, Malene Spiegelhauer, Kristina Larsen, Malene Skovsted Cilieborg, Britta Karlsson, and Jane Connie Povlsen for their assistance in animal experiments and lab analysis. The study was funded by the Novo Nordisk Foundation (DNN, Grant No. NNF220C0078747). ZY was supported by a Ph.D. grant from the China Scholarship Council while OB was supported by the BRIDGE–Translational Excellence Programme (bridge.ku.dk) at the Faculty of Health and Medical Sciences, University of Copenhagen, funded by the Novo Nordisk Foundation (Grant No. NNF23SA0087869 and NNF20SA00643). BC and TW received funding from the European Research Council under the European Union’s Horizon 2020 research and innovation programme (Grant No. 946228)

## Author contributions

DNN, OB, and ZY designed the animal experiments. OB, TW and BC planned and conducted analysis on human data. ZW and DNN performed the animal experiments. ZY performed all laboratory analyses. TM performed bacteriology and hepatic transcriptome annotation and submitted high-throughput sequence data to GEO. ZY performed all statistical analyses of clinical data, managed raw data, and generated all figures and tables. ZY, NTNT, and NTHY were responsible for omics data analysis under DNN and NPL’s supervision. ZY, OB, BK, NPL, and DNN were mainly responsible for data interpretation. ZY, OB, BK, and DNN wrote the first draft of the manuscript. All authors contributed to data interpretation, manuscript revision, and approval of the final manuscript version.

## REFERENCES

1. Stoll, B. J. et al. Trends in Care Practices, Morbidity, and Mortality of Extremely Preterm Neonates, 1993-2012. JAMA 314, 1039–1051 (2015).

2. Collins, A., Weitkamp, J. H. & Wynn, J. L. Why are preterm newborns at increased risk of infection? Arch Dis Child Fetal Neonatal Ed 103, F391–F394 (2018).

3. Harbeson, D., Francis, F., Bao, W., Amenyogbe, N. A. & Kollmann, T. R. Energy Demands of Early Life Drive a Disease Tolerant Phenotype and Dictate Outcome in Neonatal Bacterial Sepsis. Front Immunol 9, 1918 (2018).

4. Medzhitov, R., Schneider, D. S. & Soares, M. P. Disease tolerance as a defense strategy. Science 335, 936–941 (2012).

5. Wu, Z. et al. Regulation of host metabolism and defense strategies to survive neonatal infection. Biochim Biophys Acta Mol Basis Dis 1870, (2024).

6. Muk, T., Brunse, A., Henriksen, N. L., Aasmul-Olsen, K. & Nguyen, D. N. Glucose supply and glycolysis inhibition shape the clinical fate of Staphylococcus epidermidis– infected preterm newborns. JCI Insight 7, (2022).

7. Bæk, O. et al. Altered hepatic metabolism mediates sepsis preventive effects of reduced glucose supply in infected preterm newborns. Elife 13, (2024).

8. Martínez-Reyes, I. & Chandel, N. S. Mitochondrial TCA cycle metabolites control physiology and disease. Nat Commun 11, (2020).

9. Liu, M. et al. α-Ketoglutarate Modulates Macrophage Polarization Through Regulation of PPARγ Transcription and mTORC1/p70S6K Pathway to Ameliorate ALI/ARDS. Shock 53, 103–113 (2020).

10. Trauelsen, M. et al. Extracellular succinate hyperpolarizes M2 macrophages through SUCNR1/GPR91-mediated Gq signaling. Cell Rep 35, (2021).

11. Domínguez-Andrés, J. et al. The Itaconate Pathway Is a Central Regulatory Node Linking Innate Immune Tolerance and Trained Immunity. Cell Metab 29, 211–220.e5 (2019).

12. Park, J. H., Ku, H. J., Lee, J. H. & Park, J. W. Disruption of IDH2 attenuates lipopolysaccharide-induced inflammation and lung injury in an α-ketoglutarate-dependent manner. Biochem Biophys Res Commun 503, 798–802 (2018).

13. Bisgaard, H. et al. Deep phenotyping of the unselected COPSAC2010 birth cohort study. Clin Exp Allergy 43, 1384–1394 (2013).

14. Brustad, N. et al. Urban metabolic and airway immune profiles increase the risk of infections in early childhood. Thorax 79, thorax-2024-221460 (2024).

15. Thysen, A. H. et al. Distinct immune phenotypes in infants developing asthma during childhood. Sci Transl Med 12, 258 (2020).

16. Chawes, B. L. et al. Effect of Vitamin D3 Supplementation During Pregnancy on Risk of Persistent Wheeze in the Offspring: A Randomized Clinical Trial. JAMA 315, 353–361 (2016).

17. H, B., et al. Fish Oil-Derived Fatty Acids in Pregnancy and Wheeze and Asthma in Offspring. N Engl J Med 375, 81 (2016).

18. Olarini, A. et al. Vertical Transfer of Metabolites Detectable from Newborn’s Dried Blood Spot Samples Using UPLC-MS: A Chemometric Study. Metabolites 2022, Vol. 12, Page 94 12, 94 (2022).

19. Shyer, J. A., Flavell, R. A. & Bailis, W. Metabolic signaling in T cells. Cell Res 30, 649– 659 (2020).

20. Pearce, E. J. & Everts, B. Dendritic cell metabolism. Nat Rev Immunol 15, 18–29 (2015).

21. Frey, P. A. The Leloir pathway: a mechanistic imperative for three enzymes to change the stereochemical configuration of a single carbon in galactose. The FASEB Journal 10, 461– 470 (1996).

22. Strunk, T. et al. Impaired Cytokine Responses to Live Staphylococcus epidermidis in Preterm Infants Precede Gram-positive, Late-onset Sepsis. Clin Infect Dis 72, 271–278 (2021).

23. Bæk, O. et al. Diet Modulates the High Sensitivity to Systemic Infection in Newborn Preterm Pigs. Front Immunol 11, 1019 (2020).

24. Gannon, M. C., Khan, M. A. & Nuttall, F. Q. Glucose appearance rate after the ingestion of galactose. Metabolism 50, 93–98 (2001).

25. Oh, T. S., et al. Dichloroacetate improves systemic energy balance and feeding behavior during sepsis. JCI Insight 7, (2022).

26. Khaliq, W. et al. Lipid metabolic signatures deviate in sepsis survivors compared to non-survivors. Comput Struct Biotechnol J 18, 3678–3691 (2020).

27. Abramowski, A., Ward, R. & Hamdan, A. H. Neonatal Hypoglycemia. StatPearls [Internet]. Treasure Island (FL): StatPearls Publishing (2023).

28. Salis, E. R., Reith, D. M., Wheeler, B. J., Broadbent, R. S. & Medlicott, N. J. Hyperglycaemic preterm neonates exhibit insulin resistance and low insulin production. BMJ Paediatr Open 1, e000160 (2017).

29. Hofman, P. L. et al. Premature Birth and Later Insulin Resistance. New England Journal of Medicine 351, 2179–2186 (2004).

30. Zhang, W. et al. Lactate Is a Natural Suppressor of RLR Signaling by Targeting MAVS. Cell 178, 176–189.e15 (2019).

31. Conte, F., van Buuringen, N., Voermans, N. C. & Lefeber, D. J. Galactose in human metabolism, glycosylation and congenital metabolic diseases: Time for a closer look. Biochimica et Biophysica Acta (BBA) - General Subjects 1865, 129898 (2021).

32. Kalhan, S. C. et al. Estimation of gluconeogenesis in newborn infants. Am J Physiol Endocrinol Metab 281, (2001).

33. Chacko, S. K. & Sunehag, A. L. Gluconeogenesis continues in premature infants receiving total parenteral nutrition. Arch Dis Child Fetal Neonatal Ed 95, F413–F418 (2010).

34. Willmann, K. & Moita, L. F. Physiologic disruption and metabolic reprogramming in infection and sepsis. Cell Metab 36, 927–946 (2024).

35. Zhao, C. et al. Biased allosteric activation of ketone body receptor HCAR2 suppresses inflammation. Mol Cell 83, 3171–3187.e7 (2023).

36. Youm, Y. H. et al. The ketone metabolite β-hydroxybutyrate blocks NLRP3 inflammasome-mediated inflammatory disease. Nat Med 21, 263–269 (2015).

37. Brustad, N. et al. Burden of Infections in Early Life and Risk of Infections and Systemic Antibiotics Use in Childhood. JAMA Netw Open 8, e2453284 (2025).

38. Rago, D. et al. Characteristics and Mechanisms of a Sphingolipid-associated Childhood Asthma Endotype. Am J Respir Crit Care Med 203, 853–863 (2021).

39. Rago, D. et al. Fish-oil supplementation in pregnancy, child metabolomics and asthma risk. EBioMedicine 46, 399–410 (2019).

40. Sumner, L. W. et al. Proposed minimum reporting standards for chemical analysis Chemical Analysis Working Group (CAWG) Metabolomics Standards Initiative (MSI). Metabolomics 3, 211–221 (2007).

41. Brustad, N. et al. Diet-associated vertically transferred metabolites and risk of asthma, allergy, eczema, and infections in early childhood. Pediatr Allergy Immunol 34, (2023).

42. Mesotten, D. et al. ESPGHAN/ESPEN/ESPR/CSPEN guidelines on pediatric parenteral nutrition: Carbohydrates. Clinical Nutrition 37, 2337–2343 (2018).

43. Kim, D. et al. TopHat2: accurate alignment of transcriptomes in the presence of insertions, deletions and gene fusions. Genome Biol 14, R36 (2013).

44. Anders, S., Pyl, P. T. & Huber, W. HTSeq—a Python framework to work with high-throughput sequencing data. Bioinformatics 31, 166–169 (2015).

45. Love, M. I., Huber, W. & Anders, S. Moderated estimation of fold change and dispersion for RNA-seq data with DESeq2. Genome Biol 15, 550 (2014).

46. Zhu, A., Ibrahim, J. G. & Love, M. I. Heavy-tailed prior distributions for sequence count data: removing the noise and preserving large differences. Bioinformatics 35, 2084–2092 (2019).

47. Wu, T. et al. clusterProfiler 4.0: A universal enrichment tool for interpreting omics data. The Innovation 2, 100141 (2021).

48. Gu, Z., Eils, R. & Schlesner, M. Complex heatmaps reveal patterns and correlations in multidimensional genomic data. Bioinformatics 32, 2847–2849 (2016).

49. Chong, J. & Xia, J. MetaboAnalystR: an R package for flexible and reproducible analysis of metabolomics data. Bioinformatics 34, 4313–4314 (2018).

50. Bates, D., Mächler, M., Bolker, B. M. & Walker, S. C. Fitting Linear Mixed-Effects Models Using lme4. J Stat Softw 67, 1–48 (2015).

