## Supplemental Figures for "Harnessing systemic glycolysis-TCA cycle axis to boost the host defense against newborn infection"

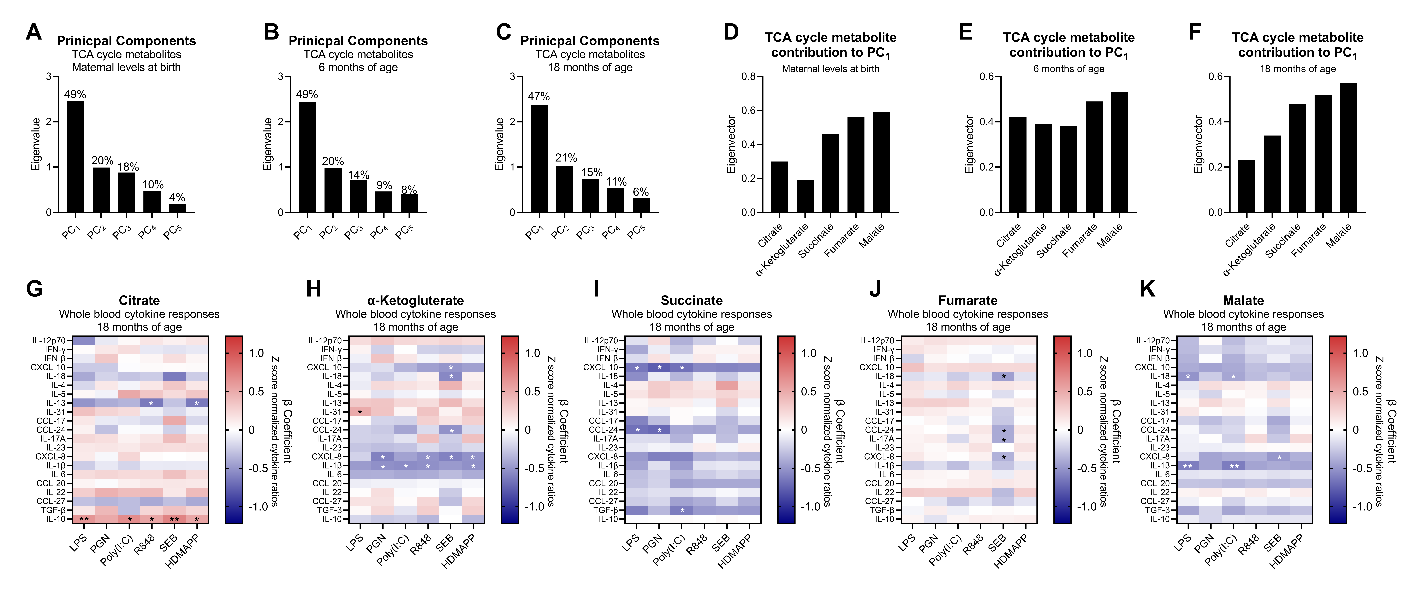

**Figure S1: Eigenvalues and eigenvectors from principal component analysis (PCAs) and the association between individual TCA cycle metabolites and immune responses at 18 months of age. (A-C):** Eigenvalues showing the fraction each principal component (PC) contributes to the overall PCA at birth, 6, and 18 months of age. **(D-F):** Eigenvectors showing how each TCA metabolite contributes to PC_1_ (TCA scores) at birth, 6, and 18 months of age. **(G-K)** Heatmaps showing the association between individual TCA cycle metabolites at 18 months of age and zscore normalized cytokine ratios following whole blood stimulation with 6 pattern recognition receptor agonists. Shown as β-coefficients following a generalized linear model where red color indicates positive, and blue negative association. LPS: Lipopolysaccharide, PGN: Peptidoglycan, R848: Imidazoquinoline, SEB Staphylococcal enterotoxin B, HDMAPP: 1-Hydroxy-2-methyl-2-buten-4-yl 4-diphosphate. *P-value < 0.05, **P-value < 0.01, and ***P-value < 0.001.

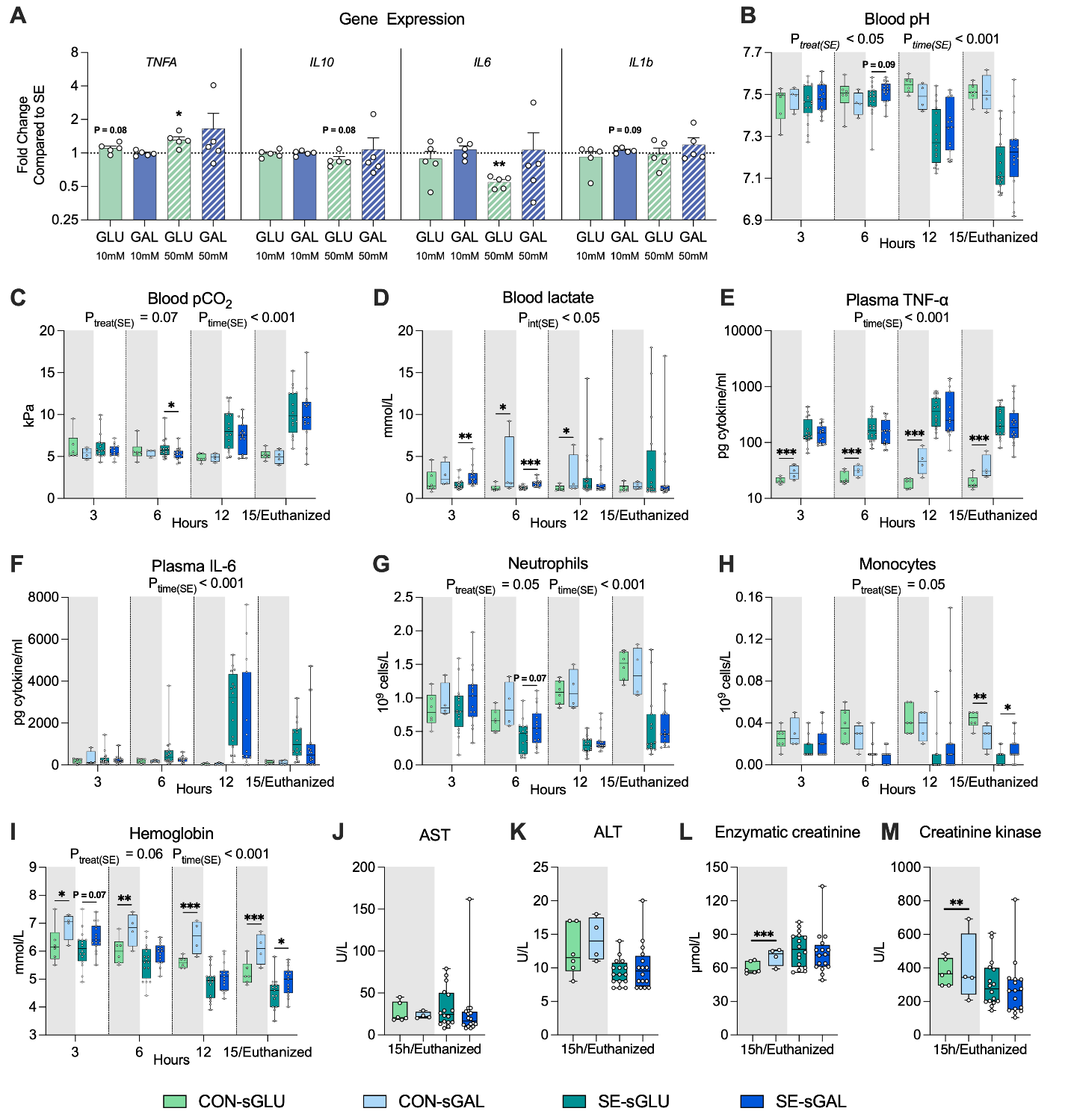

**Figure S2. Effects of galactose supply on blood gas and immune parameters during neonatal infection. (A)** Four cytokine gene expressions are shown as relative fold changes in relation to *S. epidermidis* positive control. **(B-I)** Blood and immune parameters (blood pH, pCO_2_, lactate, total leukocytes, neutrophils, monocytes, hemoglobin, plasma TNF-α and IL-6) were collected at 3, 6, 12, and 15 h post-inoculation or at euthanasia. **(J–M)** Serum biochemistry parameters (aspartate transaminase, alanine transaminase, enzymatic creatinine, and creatinine kinase) at 15 h post-inoculation or at euthanasia.  **Statistics**: Data at each time point were analyzed separately via a linear mixed-effects model, incorporating group, gender, and birth weight as fixed factors and litter as a random factor. *P-value < 0.05, **P-value < 0.01, and ***P-value < 0.001, compared between SE-sGAL and SE-sGLU groups at the same time point. Another linear mixed-effects model was employed to probe further disparities spanning the entire experimental duration, incorporating group, time, their interaction, gender, and birth weight as fixed factors, with litter and pig ID as random factors. P_treat(SE)_, P_time(SE)_, and P_int(SE)_ denote probability values for group effect (SE-sGAL and SE-sGLU) over time, time effects, and the interaction effects between time and group in the linear mixed effects interaction model, respectively. Uninfected animals (CON) served as a reference group and were not compared directly with infected animals (SE). Statistical significance was defined as P-value < 0.05. All data are presented as box and whisker plots showing the range from minimum to maximum values.

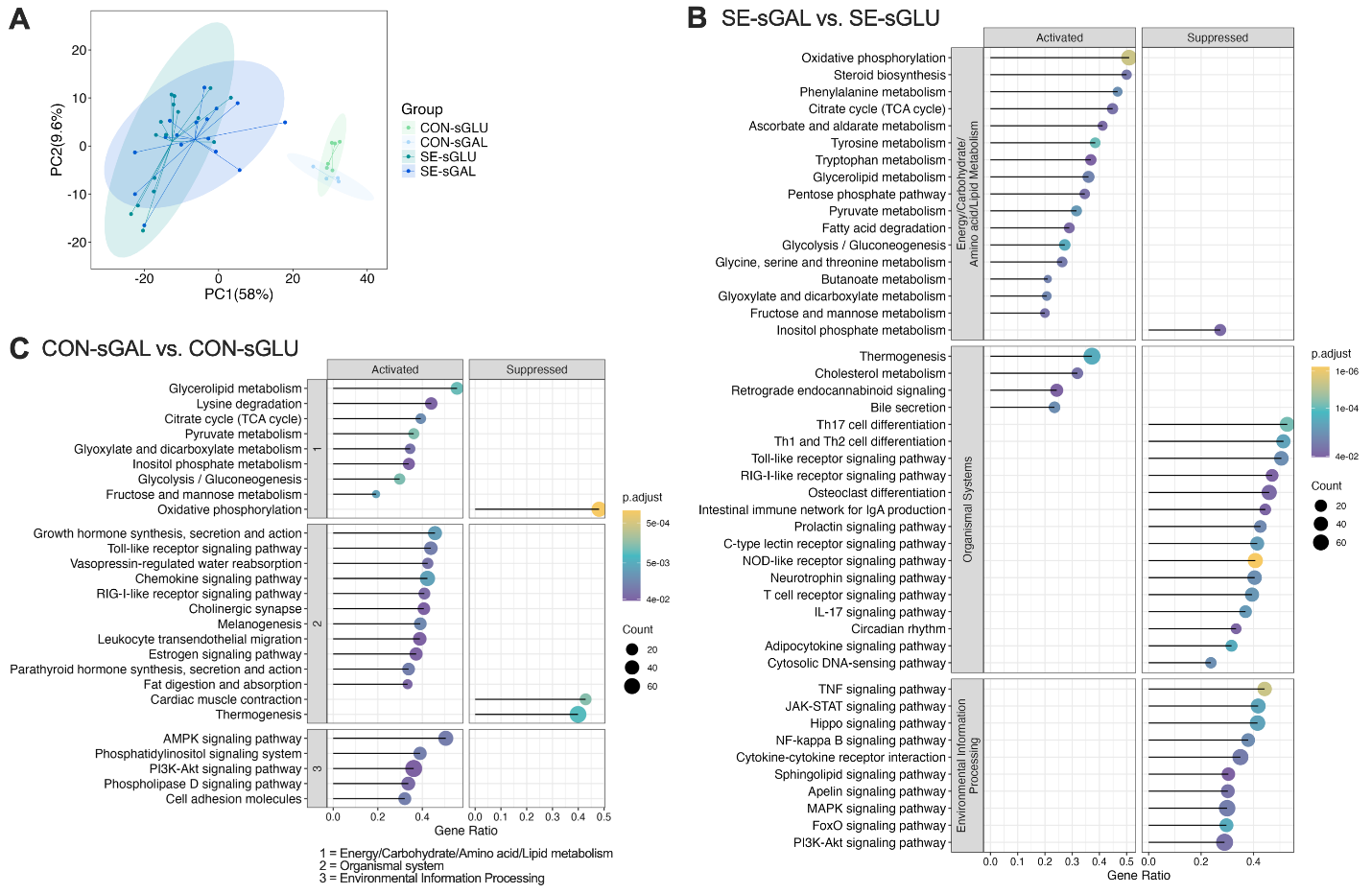

**Figure S3. Galactose supply reduces inflammation and enhances multiple hepatic metabolic pathways. (A)** PCA scores plot of the first two principal components. **(B,C)** GSEA was performed between the SE-sGAL vs SE-sGLU as well as infected vs uninfected groups, respectively. The *Sus scrofa* (pig) KEGG knowledgebase was utilized for pathway enrichment analysis. Significant pathways in specific categories such as energy metabolism, carbohydrate metabolism, amino acid metabolism, lipid metabolism, signal transduction, and immune system have been chosen for presentation. The complete list of enriched pathways can be found in **Table S1A**. The size and color of the dots indicate the gene ratio and FDR values, respectively.

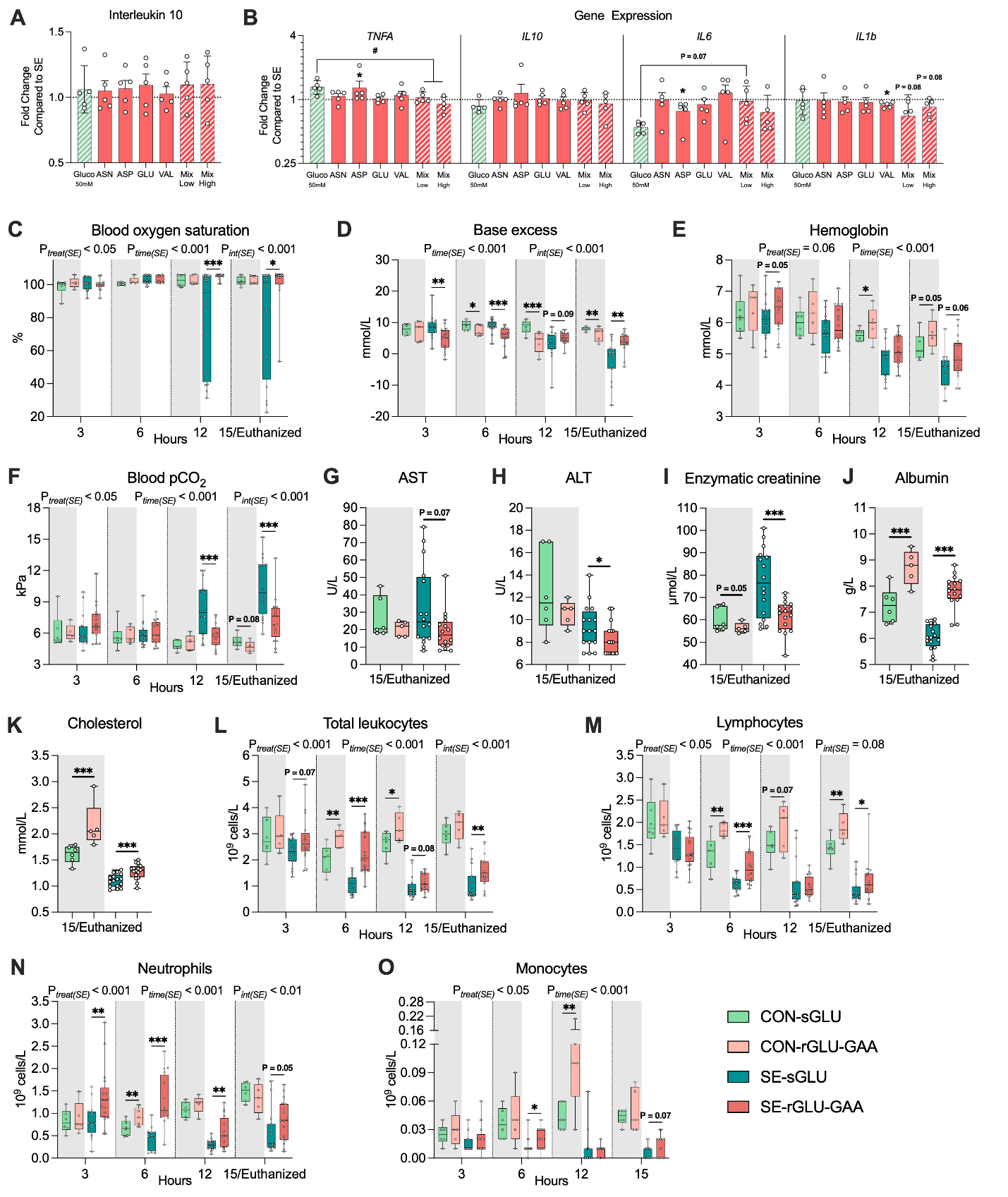

**Figure S4. Combined glucose restriction and glucogenic amino acid supply enhances both, host disease resistance and tolerance. (A)** *In vitro* human macrophage-like THP1 cells exposed to live *S. epidermidis*. Levels of interleukin 10 are shown as relative fold changes in relation to *S. epidermidis* positive control. **(B)** Four cytokine gene expressions are shown as relative fold changes in relation to *S. epidermidis* positive control. **(C–F)** Blood pCO_2_, oxygen saturation, base excess, and hemoglobin at 3, 6, 12, and 15 h post-bacterial inoculation or at euthanasia. **(G–K)** Serum aspartate transaminase (AST), alanine transaminase (ALT), enzymatic creatine, and albumin at 15 h post-inoculation or at euthanasia. **(L–O)** Blood immune parameters (blood total leukocytes, lymphocytes, neutrophils, and monocytes) at 3, 6, 12, and 15 h post-bacterial inoculation or at euthanasia. **Statistics:** **(C–O)** Data at each time point were analyzed using a linear mixed-effects model, incorporating group, gender, and birth weight as fixed factors and litter as a random factor. *P-value < 0.05, **P-value < 0.01, and ***P-value < 0.001, compared between SE-rGLU-GAAs and SE-sGLU groups at the same time point. **(C–F & L–O)** Another linear mixed-effects model was employed to probe further disparities spanning the entire experimental duration, incorporating group, time, their interaction, gender, and birth weight as fixed factors, with litter and pig ID as random factors. P_treat(SE)_, P_time(SE)_, and P_int(SE)_ denote probability values for group effect (SE-rGLU-GAAs and SE-sGLU) over time, time effects, and the interaction effects between time and group in the linear mixed effects interaction model, respectively. Uninfected animals (CON) served as a reference group and were not compared directly with infected animals (SE). Statistical significance was defined as P-value < 0.05. All data were presented as box and whisker plots showing the range from minimum to maximum values.

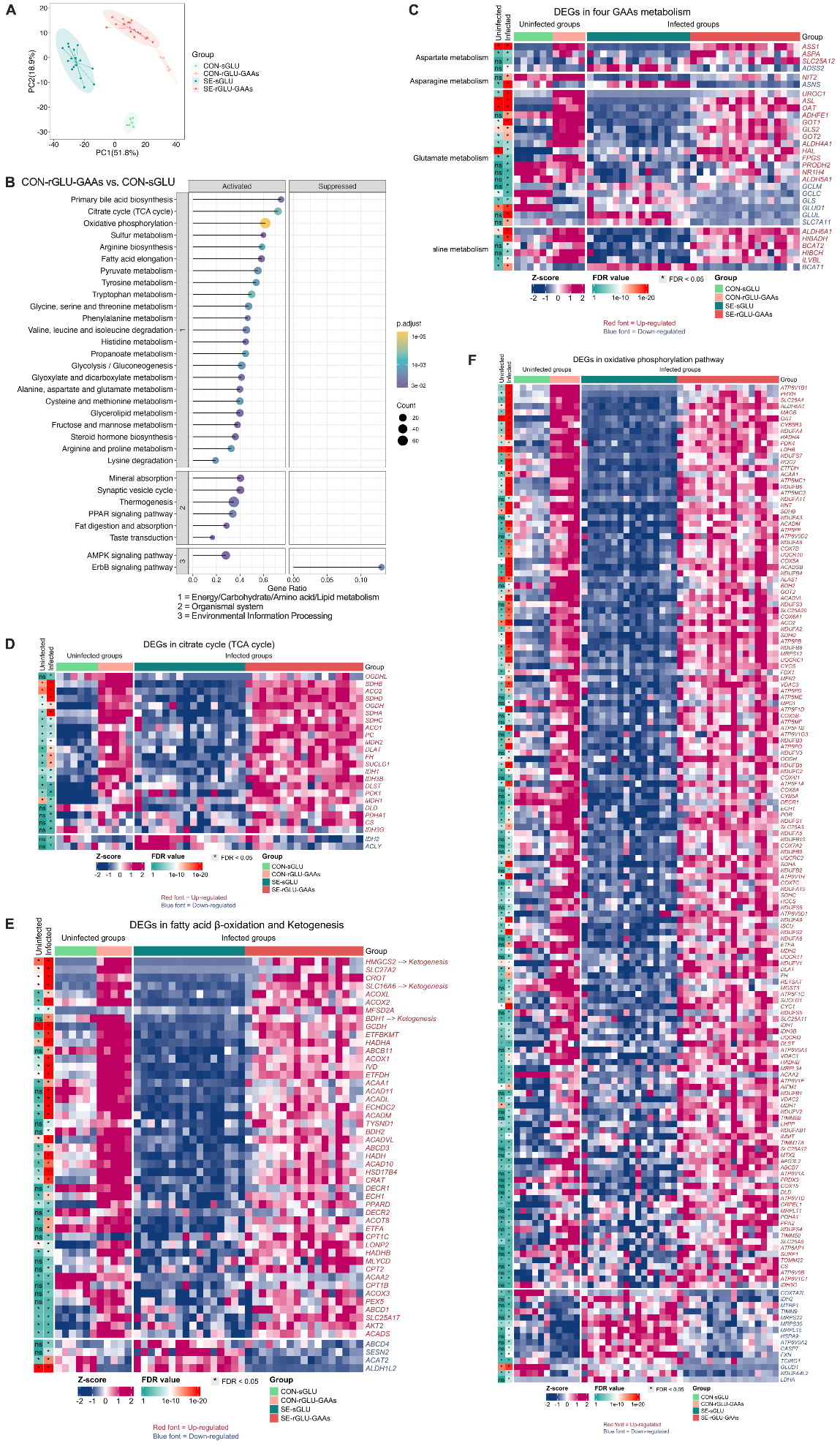

**Figure S5. Effects of rGLU-GAAs supply on hepatic metabolism. (A)** PCA of hepatic transcriptome between all groups **(B)** GSEA was performed between the CON-rGLU-GAAs and CON-sGLU groups using the *Sus scrofa* (pig) KEGG knowledgebase. Significant pathways in specific categories such as energy metabolism, carbohydrate metabolism, amino acid metabolism, lipid metabolism, signal transduction, and immune system have been chosen for presentation. The complete list of enriched pathways can be found in **Table S2B**. The size and color of the dots indicate the gene ratio and FDR values, respectively. **(C–F)** Heatmaps illustrating DEGs involved in the specific metabolic pathways between SE-rGLU-GAAs and SE-sGLU groups.

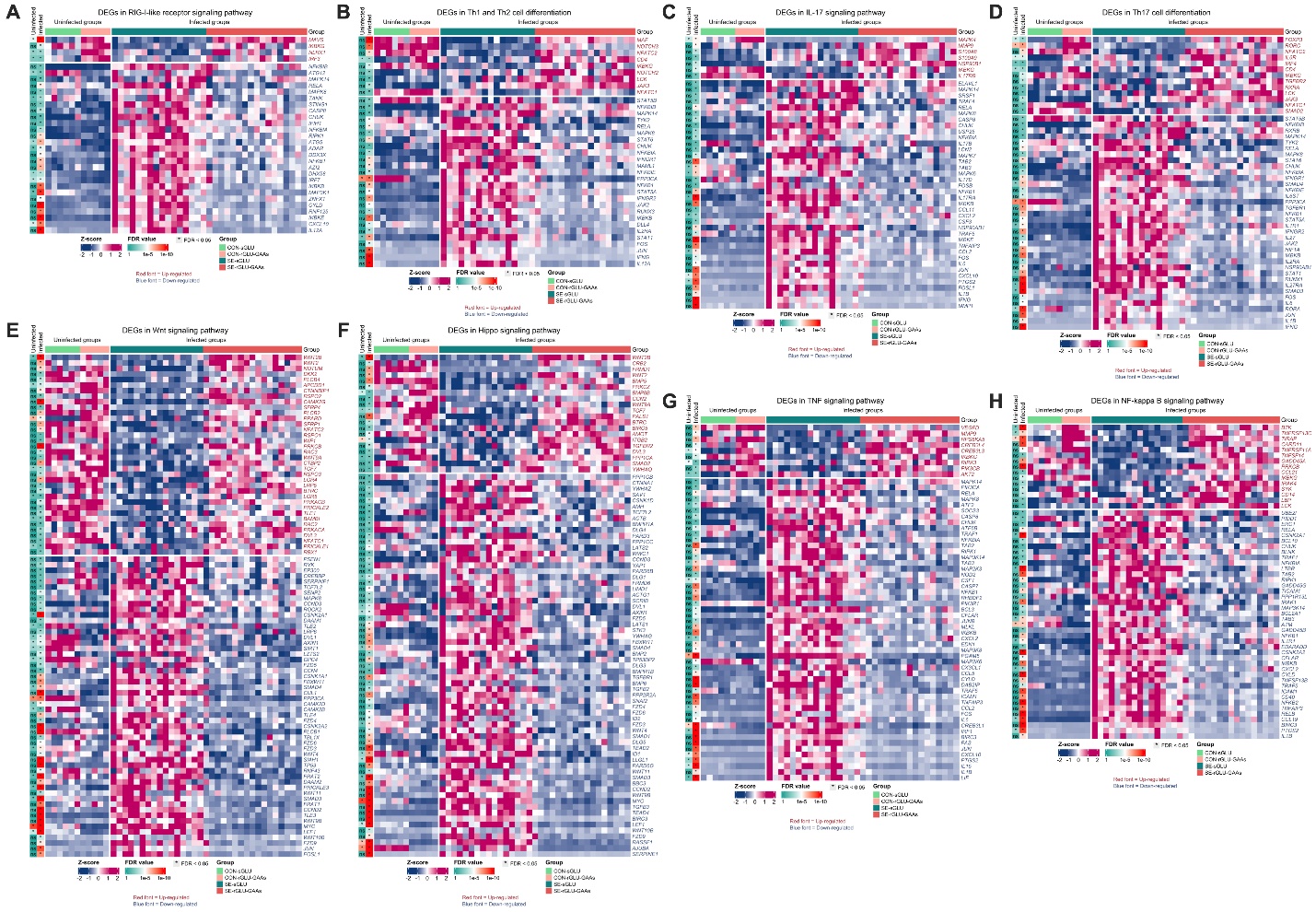

**Figure S6. rGLU-GAAs supply suppressed hepatic pathways related to immunity and signal transduction. (A-H)** Heatmaps illustrating DEGs involved in the critical immune/inflammation-related pathways between SE-rGLU-GAAs and SE-sGLU groups.

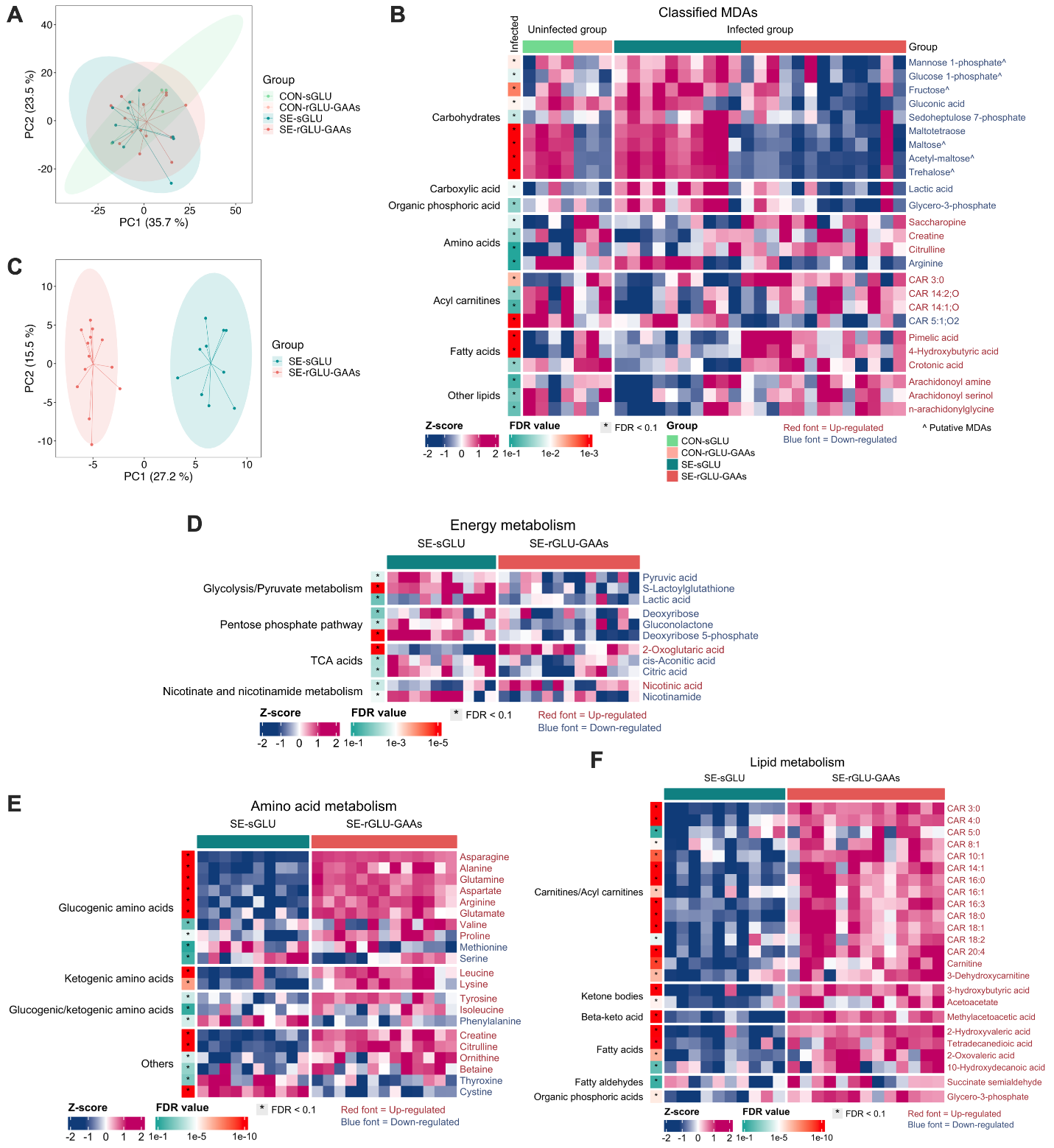

**Figure S7. rGLU-GAAs supply during infection affect plasma and hepatic metabolome. (A-B)** PCA analysis and heatmap showing metabolites with differential abundances from hepatic metabolome. (C-E) PCA analysis and heatmaps showing metabolites with differential abundances in different metabolic categories.

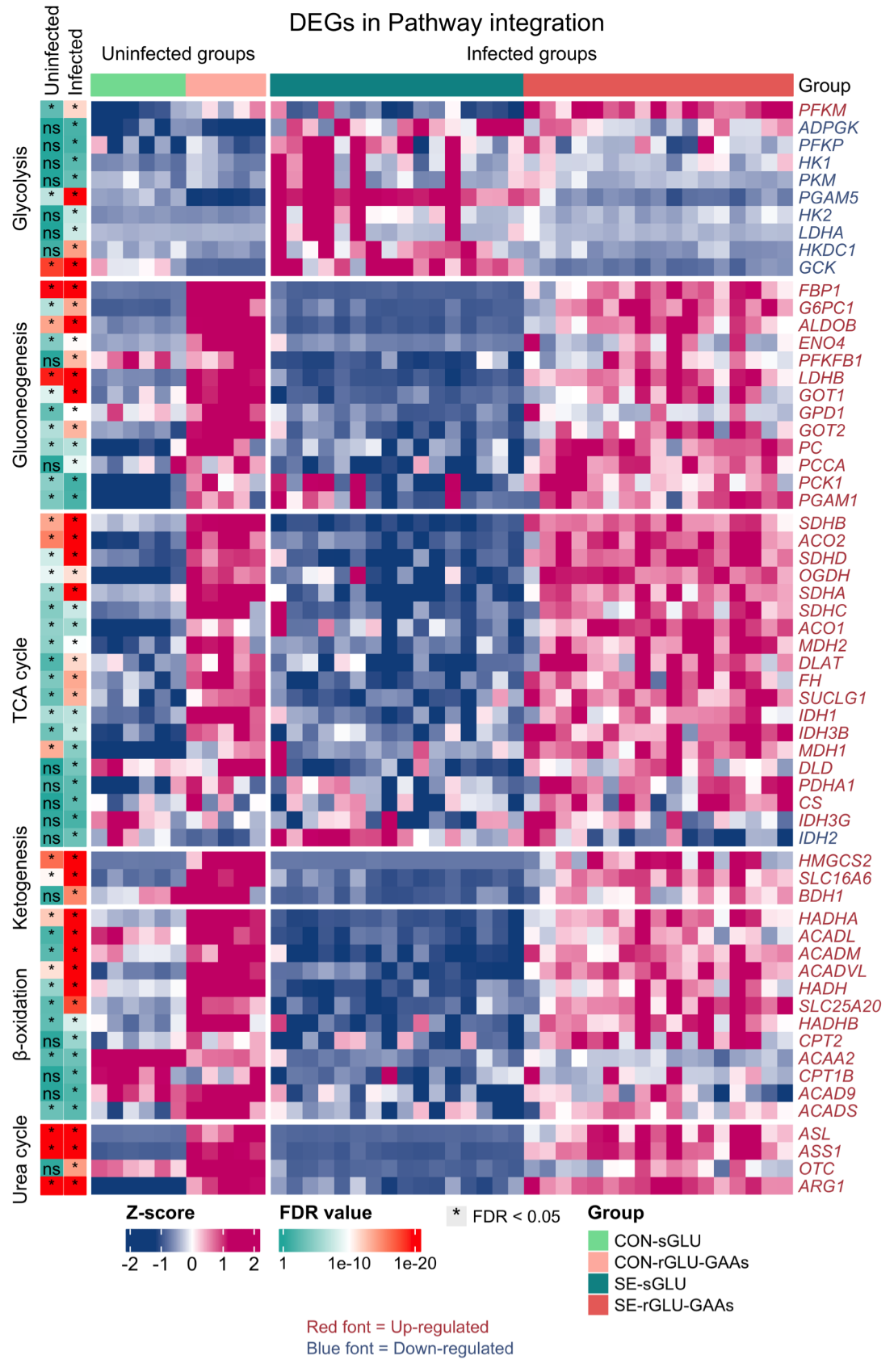

**Figure S8. Effects of rGLU-GAAs supply on hepatic metabolism. (A-H)** Heatmap illustrating DEGs between SE-rGLU-GAAs and SE-sGLU groups involved in **Figure 5F**.

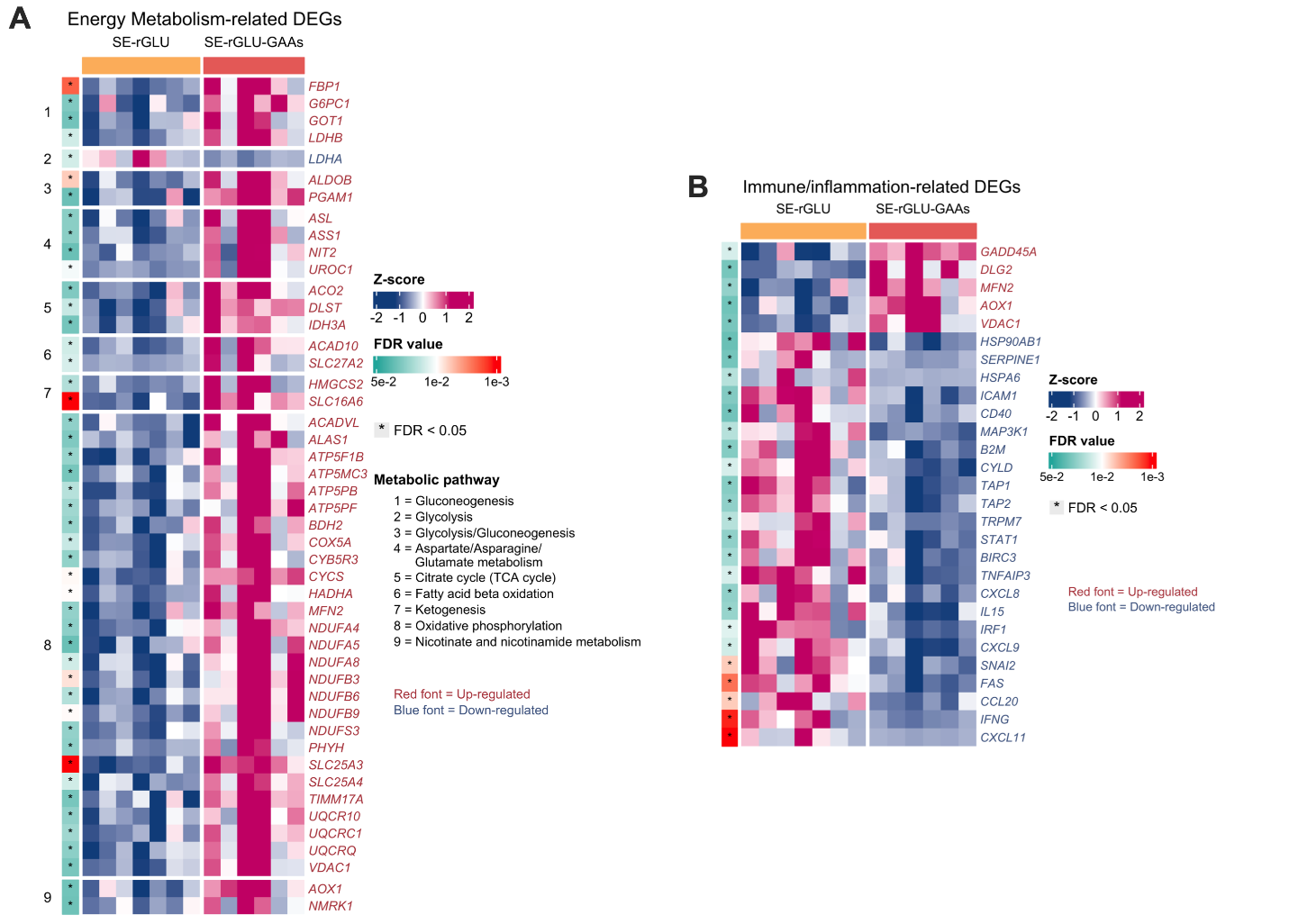

**Figure S9. Effects of GAA supply during glucose restriction on hepatic gene expressions related to energy metabolism (A) and inflammation (B)**.
