## Supplemental Table S1 for "Harnessing systemic glycolysis-TCA cycle axis to boost the host defense against newborn infection"

**Supplementary Table 1: Infection rate ratios and hazard ratios for each individual TCA cycle metabolite**

|  | **Citrate** | | **α-Ketogluterate** | | | **Succinate** | | | **Fumarate** | | **Malate** | |
| --- | --- | --- | --- | --- | --- | --- | --- | --- | --- | --- | --- | --- |
| **Infection risk**  **Birth to 36 months** | **IRR (95% CI)** | **P** | **IRR (95% CI)** | | **P** | **IRR (95% CI)** | | **P** | **IRR (95% CI)** | **P** | **IRR (95% CI)** | **P** |
| All infections | 0.87 (0.62 - 1.24) | 0.45 | 1.23 (0.88 - 1.44) | | 0.36 | 0.78 (0.56 - 1.09) | | 0.15 | **0.71 (0.52 - 0.96)** | **0.03** | **0.71 (0.52 - 0.96)** | **0.03** |
| Pneumonia | 0.51 (0.15 - 1.77) | 0.29 | 1.70 (0.75 - 3.88) | | 0.21 | 0.42 (0.11 - 1.53) | | 0.19 | 0.85 (0.18 - 3.99) | 0.83 | 0.81 (0.23 - 2.92) | 0.75 |
| Acute Otitis Media | 0.78 (0.27 - 2.21) | 0.63 | 0.67 (0.33 - 1.33) | | 0.25 | 0.92 (0.39 - 2.21) | | 0.86 | 0.71 (0.28 - 1.81) | 0.47 | 0.98 (0.42 - 2.29) | 0.96 |
| Tonsilltis | 1.62 (0.44 - 5.90) | 0.47 | 0.53 (0.20 - 1.40) | | 0.20 | 0.41 (0.08 - 2.12) | | 0.29 | **0.28 (0.08 - 0.93)** | **0.04** | **0.27 (0.09 - 0.80)** | **0.02** |
| Fever | 1.04 (0.65 - 1.66) | 0.88 | **1.38 (1.01 - 1.89)** | | **0.05** | 0.73 (0.45 - 1.19) | | 0.20 | **0.65 (0.42 - 0.99)** | **0.04** | 0.74 (0.49 - 1.12) | 0.15 |
| Cold symptoms | 0.70 (0.44 - 1.12) | 0.14 | 1.12 (0.80 - 1.56) | | 0.50 | 0.72 (0.47 - 1.10) | | 0.13 | **0.64 (0.43 - 0.95)** | **0.03** | **0.61 (0.42 - 0.90)** | **0.01** |
| Gastroenteritis | 1.29 (0.65 - 2.58) | 0.47 | 0.98 (0.62 - 1.55) | | 0.94 | 0.82 (0.36 - 1.90) | | 0.65 | 0.66 (0.32 - 1.32) | 0.24 | 0.77 (0.39 - 1.51) | 0.45 |
| **Infection risk**  **18 to 36 months** | **IRR (95% CI)** | **P** | **IRR (95% CI)** | | **P** | **IRR (95% CI)** | | **P** | **IRR (95% CI)** | **P** | **IRR (95% CI)** | **P** |
| All infections | **0.79 (0.66 - 0.95)** | **0.01** | 0.94 (0.80 - 1.10) | | 0.44 | 0.93 (0.75 - 1.14) | | 0.47 | **0.80 (0.72 - 0.90)** | **0.00** | 1.06 (0.91 - 1.23) | 0.44 |
| Pneumonia | 1.37 (0.49 - 3.86) | 0.55 | 0.66 (0.25 - 1.72) | | 0.40 | 1.53 (0.55 - 4.25) | | 0.41 | 1.00 (0.51 - 1.95) | 1.00 | 1.39 (0.60 - 3.24) | 0.44 |
| Acute Otitis Media | 0.89 (0.42 - 1.86) | 0.75 | 0.73 (0.38 - 1.42) | | 0.36 | 0.64 (0.26 - 1.56) | | 0.32 | **0.64 (0.41 – 1.00)** | **0.05** | 0.76 (0.41 - 1.40) | 0.38 |
| Tonsilltis | 0.97 (0.25 - 3.70) | 0.97 | 1.80 (0.58 - 5.57) | | 0.31 | 2.35 (0.77 - 7.21) | | 0.14 | 0.56 (0.25 - 1.25) | 0.16 | 2.35 (0.82 - 6.76) | 0.11 |
| Fever | 0.85 (0.59 - 1.21) | 0.36 | 1.18 (0.87 - 1.60) | | 0.30 | 0.88 (0.59 - 1.33) | | 0.56 | 0.95 (0.76 - 1.19) | 0.68 | 1.29 (0.97 - 1.73) | 0.08 |
| Cold symptoms | **0.77 (0.60 - 0.98)** | **0.03** | 0.88 (0.71 - 1.09) | | 0.26 | 0.96 (0.73 - 1.27) | | 0.78 | **0.77 (0.66 - 0.90)** | **0.00** | 1.03 (0.84 - 1.26) | 0.77 |
| Gastroenteritis | 0.50 (0.26 - 0.97) | 0.04 | 0.78 (0.44 - 1.39) | | 0.40 | 0.63 (0.29 - 1.39) | | 0.25 | 0.74 (0.49 - 1.10) | 0.14 | 0.64 (0.38 - 1.10) | 0.10 |
| **Time to first infection**  **Birth to 36 months** | **HR (95% CI)** | **P** | **HR (95% CI)** | | **P** | **HR (95% CI)** | | **P** | **HR (95% CI)** | **P** | **HR (95% CI)** | **P** |
| All infections | 0.97 (0.46-2.03) | 0.93 | 1.60 (0.95-2.69) | | 0.08 | 0.64 (0.32-1.28) | | 0.21 | 0.75 (0.36-1.55) | 0.44 | 0.83 (0.43-1.59) | 0.58 |
| Pneumonia | 0.37 (0.10-1.32) | 0.13 | 2.05 (0.96-4.40) | | 0.07 | 0.25 (0.07-0.90) | | 0.03 | 0.39 (0.12-1.27) | 0.12 | 0.42 (0.14-1.23) | 0.12 |
| Acute Otitis Media | 0.86 (0.34-2.18) | 0.74 | 1.12 (0.60-2.11) | | 0.73 | 0.69 (0.28-1.73) | | 0.43 | 0.45 (0.19-1.09) | 0.08 | 0.66 (0.29-1.52) | 0.33 |
| Tonsilltis | 1.91 (0.49-7.53) | 0.35 | 0.37 (0.13-1.03) | | 0.06 | 0.45 (0.10-1.91) | | 0.28 | **0.22 (0.06-0.86)** | **0.03** | **0.20 (0.06-0.71)** | **0.01** |
| Fever | 0.91 (0.44-0.90) | 0.80 | **2.15 (1.27-3.65)** | | **<0.01** | 1.19 (0.57-2.47) | | 0.64 | 1.25 (0 .58-2.70) | 0.56 | 1.78 (0.89-3.54) | 0.10 |
| Cold symptoms | 0.90 (0.43-1.87) | 0.77 | 1.41 (0.84-2.36) | | 0.19 | 0.82 (0.40-1-66) | | 0.58 | 0.88 (0.44-1.78) | 0.73 | 0.85 (0.45-1.63) | 0.63 |
| Gastroenteritis | 1.43 (0.62-3.31) | 0.39 | 0.89 (0.52-1.54) | | 0.68 | 0.71 (0.34-1.51) | | 0.37 | 0.87 (0.40-1.89) | 0.72 | 1.30 (0.66-2.59) | 0.45 |
| **Time to first infection**  **18 to 36 months** | **HR (95% CI)** | **P** | **HR (95% CI)** | | **P** | **HR (95% CI)** | | **P** | **HR (95% CI)** | **P** | **HR (95% CI)** | **P** |
| Pneumonia | 1.14 (0.17-7.43) | 0.89 | 0.69 (0.10-4.82) | | 0.70 | 0.23 (0.01-3.80) | | 0.31 | 0.50 (0.12-2.19) | 0.36 | 0.73 (0.13-4.22) | 0.72 |
| Acute Otitis Media | 0.57 (0.13-2.64) | 0.48 | 0.71 (0.16-3.19) | | 0.66 | 0.36 (0.04-3.10) | | 0.35 | 0.30 (0.07-1.03) | 0.06 | 0.80 (0.20-3.22) | 0.75 |
| Tonsilltis | 0.53 (0.06-4.66) | 0.57 | 1.88 (0.27-13.1) | | 0.52 | 7.72 (0.93-63.9) | | 0.06 | 0.54 (0.11-2.84) | 0.48 | 3.93 (0.58-26.9) | 0.16 |
| Fever | 0.56 (0.21-1.50) | 0.25 | 1.47 (0.58-3.70) | | 0.41 | 0.58 (0.17-2.01) | | 0.39 | 0.84 (0.41-1.74) | 0.64 | 1.14 (0.49-2.68) | 0.76 |
| Cold symptoms | 1.04 (0.37-2.93) | 0.94 | 1.17 (0.45-3.04) | | 0.75 | 1.23 (0.36-4.20) | | 0.74 | 0.94 (0.42-2.11) | 0.88 | 0.95 (0.37-2.42) | 0.91 |
| Gastroenteritis | 0.34 (0.09-1.32) | 0.12 | 1.28 (0.35-4.76) | 0.71 | | 0.24 (0.04-1.59) | 0.14 | | 1.69 (0.56-5.11) | 0.35 | 0.85 (0.25-2-89) | 0.79 |

Association between plasma relative abundance levels of TCA cycle metabolites and early life infections. Infection risk estimated by quasi-poission regression and time to first infection estimated by a cox proportional hazard model. Both shown from birth until 36 months of age and from 18 to 36 months of age. CI: confidence interval, IRR: Infection Rate Ratio, HR: Hazard Ratio.
